# The Centralspindlin proteins Pavarotti and Tumbleweed work with WASH to regulate Nuclear Envelope budding

**DOI:** 10.1101/2022.11.16.516846

**Authors:** Kerri A. Davidson, Mitsutoshi Nakamura, Jeffrey M. Verboon, Susan M. Parkhurst

**Author notes:** These authors contributed equally (Kerri A. Davidson, Mitsutoshi Nakamura). Corresponding author: Fred Hutchinson Cancer Center, 1100 Fairview Ave N., A1-187, Seattle, WA, 98109, USA.

## Abstract

Nuclear envelope (NE) budding is a nuclear pore independent nuclear export pathway, analogous to the egress of herpesviruses, and required for protein quality control, synapse development and mitochondrial integrity. The physical formation of NE buds is dependent on the Wiskott-Aldrich Syndrome protein Wash, its regulatory complex (SHRC), and Arp2/3, and requires Wash’s actin nucleation activity. However, the machinery governing cargo recruitment and organization within the NE bud remains unknown. Here, we identify Pavarotti (Pav) and Tumbleweed (Tum) as new molecular components of NE budding. Pav and Tum interact directly with Wash and define a second nuclear Wash-containing complex required for NE budding. Interestingly, we find that the actin bundling activities of Wash and Pav are required, suggesting a structural role in the physical and/or organizational aspects of NE buds. Thus, Pav and Tum are providing exciting new entry points into the physical machineries of this alternative nuclear export pathway for large cargos during cell differentiation and development.

## INTRODUCTION

Trafficking of macromolecules from the nucleus to the cytoplasm is a major cellular process that is critical for all developmental processes, including aging, neuron development, and regulation of diferentiation, and when dysregulated, is associated with diseases and cancers (Burke and Stewart, 2014; Grunwald et al., 2011; Siddiqui and Borden, 2012; Tran et al., 2014). Nuclear Envelope (NE) budding is an alternative nuclear export pathway for large macromolecular machineries and complexes, including large ribonucleoprotein (megaRNP) granules (Speese et al., 2012). These megaRNPs can be preassembled to deliver and allow for co-regulation of functionally-related components required for major developmental pathways or to remove obsolete protein aggregates all while bypassing canonical nuclear export through nuclear pores (Fradkin and Budnik, 2016; Hatch and Hetzer, 2014; Jokhi et al., 2013; Li et al., 2016; Panagaki et al., 2021; Parchure et al., 2017; Ramaswami et al., 2013; Rose and Schlieker, 2012; Speese et al., 2012; Verboon et al., 2020). During NE budding export, megaRNPs or other cargoes are encircled by the nuclear lamina (including type-A and -B lamins) and the inner nuclear membrane (INM). This initial bud is then pinched off from the INM and subsequently fuses with the outer nuclear membrane (OMN) to release the megaRNP/cargo into the cytosplasm (Fig. 1A-B).

**Fig. 1.**
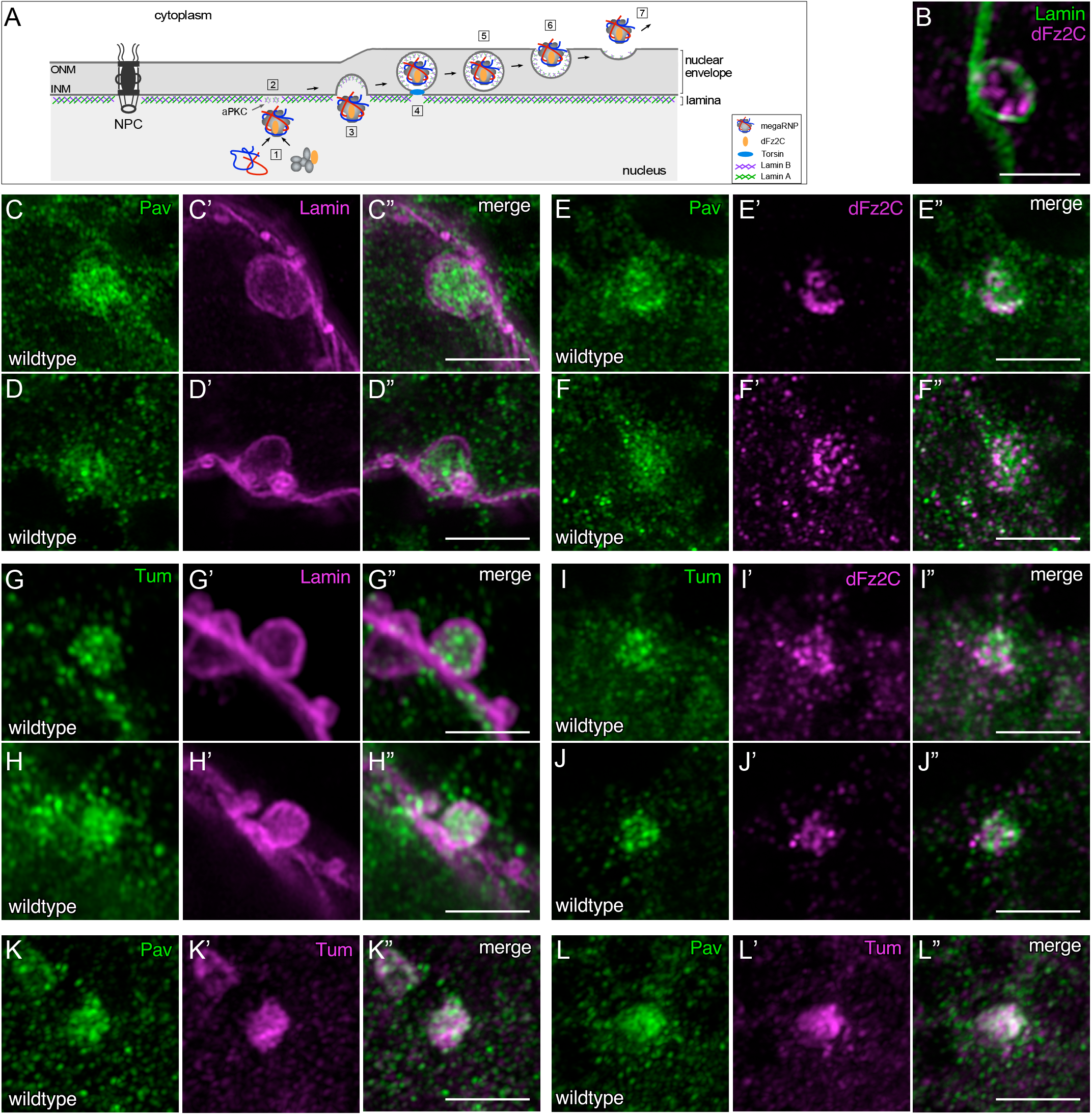
Pav and Tum are expressed in the nucleus and accumulate at NE-buds. (A) Schematic of NE-budding steps: megaRNPs visualized by dFz2C are assembled [1], the nuclear lamina is modified by aPKC [2], the inner nuclear membrane (INM) and lamina begins to bend and megaRNPs enter this membrane deformation [3] and subsequently are encapsulated by the INM [4], scission of the INM occurs [5], followed by INM fusion with the outer nuclear membrane (ONM) [6], and megaRNP exit into cytoplasm [7]. (B) Super-resolution (Airyscan) micrograph projection of wildtype larval salivary gland nucleus stained with antibodies to Lamin B and dFz2C. Large Lamin B and dFz2C positive puncta indicate NE-buds. (C-L”) Super-resolution micrograph projections of NE buds from wildtype larval salivary gland nuclei stained with antibodies to Lamin B and Pav (C-D”), Pav and dFz2C (E-F”), Lamin B and Tum (G-H”), Tum and dFz2C (I-J”), and Pav and Tum (K-L”). Scale bars: 0.5μm.

NE budding is an evolutionarily conserved process: endogenous perinuclear foci/buds have been observed in a variety of plant and animal cells (Dickinson and Potter, 1975; Dickinson and Potter, 1979; Hadek and Swift, 1962; LaMassa et al., 2018; Panagaki et al., 2018; Szollosi, 1965; Szollosi and Szollosi, 1988). NE budding has been shown to be important for neuro-muscular junction integrity affecting synapse development, mitochondrial integrity leading to premature aging in *Drosophila*, and for the removal of aggregated proteins following cellular stress in *Saccharomyces cerevisiae* (Jokhi et al., 2013; Li et al., 2016; Panagaki et al., 2021; Speese et al., 2012). However, the full spectrum of biological processes or contexts in which megaRNPs or other large macromolecular cargos require and/or utilize the NE budding pathway is just beginning to be known. NE budding also shares many similarities with nuclear egress mechanisms described for herpesviruses, widespread pathogens that cause or contribute to a diverse array of human diseases (Bigalke and Heldwein, 2015; Bigalke and Heldwein, 2016; Johnson and Baines, 2011; Lee and Chen, 2010; Mettenleiter et al., 2013; Parchure et al., 2017).

We recently identified Wash, a Wiskott-Aldrich Syndrome (WAS) family protein, and its four subunit WASH Regulatory Complex (SHRC; consisting of CCDC53, Strumpellin, SWIP and FAM21) as new molecular components involved in the physical aspects of NE bud formation (Verboon et al., 2020). WAS family proteins polymerize branched actin filaments through the Arp2/3 complex, and are often involved in membrane-cytoskeletal interactions and organization, including membrane deformations needed for endocytosis or formation of cellular protrusions (Takenawa and Suetsugu, 2007). We found that Wash is involved in two nuclear functions that affect NE budding: (1) Wash <u>indirectly</u> affects NE budding through its physical interaction with Lamin B leading to general nuclear lamina disruption and inefficient NE bud formation. This nonspecific Wash activity is SHRC- and Arp2/3- <u>in</u>dependent. (2) Wash <u>directly</u> affects NE budding. Wash, along with its SHRC, Arp2/3, and capping protein, is needed for the physical formation of NE buds (Verboon et al., 2020).

The *Drosophila* MKLP1 ortholog Pavarotti (Pav), is a kinesin-like protein that, along with Tumbleweed (Tum; MgcRacGAP), forms the centralspindlin complex, which bundles and moves along microtubules during cytokinesis (D’Avino et al., 2015; Green et al., 2012; Mishima et al., 2002; Pollard and O’Shaughnessy, 2019; Somers and Saint, 2003; White and Glotzer, 2012). Our recent work has shown that Pav, but not Tum, can also bind directly to F-actin, and that this activity is needed for its roles in cell wound repair and oogenesis (Nakamura et al., 2020). In addition to their roles in cytoplasm, Tum and Pav function in the nucleus to regulate Wnt signaling (Jones et al., 2010). Here we show that Pav and Tum can bind directly to Wash, and are required for NE budding. Pav and Tum proteins are both enriched in NE buds. Pav and Tum knockdowns lack NE buds and display phenotypes associated with the loss of NE budding. Wash is present in several independent nuclear complexes. Interestingly, we find that only Tum is associated with the Wash nuclear complex (also containing Wash’s four subunit SHRC) previously associated with NE budding, while Pav and Tum are both present in a separate Wash nuclear complex (lacking the SHRC). Thus, Wash appears to function directly in NE budding through at least two independent nuclear complexes. We also find that Wash and Pav actin bundling activities are required for proper NE budding. We propose that along with Wash, Pav and Tum play essential roles in the physical/structural aspects of NE buds.

## RESULTS

### Pav and Tum accumulate in NE buds in *Drosophila* salivary gland nuclei

To identify nuclear Wash-interacting proteins, and in particular, those working with Wash/SHRC in NE budding, we performed a mass spectometry screen with Tap-tagged Wash in *Drosophila* Kc cell nuclear lysate and a pilot proximity proteomics screen with the NE bud megaRNP cargo dFz2C in *Drosophila* S2 cells (JMV and SMP, screens to be described elsewhere). Strikingly, one of the overlapping hits from these screens was the kinesin-like protein Pavarotti (Pav), suggesting that Pav interacts with Wash and is present in NE buds. We stained larval salivary glands with an antibody specific to Pav, along with antibodies to either Lamin B or dFz2C (Fig. 1C-F”; Fig. S1). While Pav is present in the cytoplasm and nucleus, it is enriched in lamin-encircled NE buds (Fig. 1C-D”; Fig. S1A-A”). Pav exhibits a punctate distribution within NE buds and weaves around the dFz2C puncta present within NE buds (Fig. 1E-F”), supporting a role for Pav in NE budding.

Pav was first characterized for its role in organizing the central spindle and contractile ring during cytokinesis, where it works with the Rho GTPase activating protein, Tumbleweed (Tum; RacGAP50C), to form the heterotetrameric centralspindlin complex (Adams et al., 1998; Somers and Saint, 2003; Zavortink et al., 2005). To determine whether Tum is also involved in NE budding, we stained larval salivary glands with antibody specific to Tum, along with antibodies to either Lamin B or dFz2C (Fig. 1G-J”). We find that Tum is also enriched in lamin encircled NE buds (Fig. 1G-H”), and weaves around the dFz2C puncta within the NE buds (Fig. 1I-J”), supporting a role for Tum in NE budding.

We next co-stained Pav and Tum and observe an incomplete overlap of the two proteins within NE buds (Fig. 1K-L”). Pav and Tum work together as the centralspindlin complex during cytokinesis, but they show separate and distinct distributions during single cell wound repair and function independently during *Drosophila* oogenesis (Nakamura et al., 2020). As such, the partial co-localizationof Pav and Tum within NE buds is unable to inform whether they are working together as a complex, have independent roles, a combination of both, or are merely packaged as cargos in the process of NE budding.

### Pav and Tum are required for NE bud formation and knockdowns exhibit phenotypes associated with loss of NE budding

To further elucidate the presence of Pav and Tum in NE buds, we co-labeled Pav or Tum knockdown larval salivary gland nuclei for Lamin B and dFz2C. We generated the Pav or Tum knockdowns by expressing RNAi constructs (two independent RNAi constructs for each) specifically in the larval salivary gland using the GAL4-UAS system (Fig. 2A-D; Fig. S2A-B”). We find that Pav knockdowns exhibit an average of 0.29±0.1 (n=90) and 0.15±0.0 (n=125) dFz2C foci/NE-buds respectively, compared to an average of 6.66±0.2 dFz2C foci/NE buds in control (+/Sgs-Gal4); n=125, p<0.0001) (Fig. 2A-B”, D; Fig. S2A-A”). Similarly, we find that Tum knockdowns exhibit an average of 0.24±0.1 (n=91) and 0.94±0.1 (n= 105) dFz2C foci/NE buds, respectively (Fig. 2C-D; Fig. S2B-B”). These reductions in dFz2C foci/NE buds are similar to those we reported previously for knockdowns of Wash or its SHRC in larval salivary gland nuclei (Verboon et al., 2020).

**Fig. 2.**
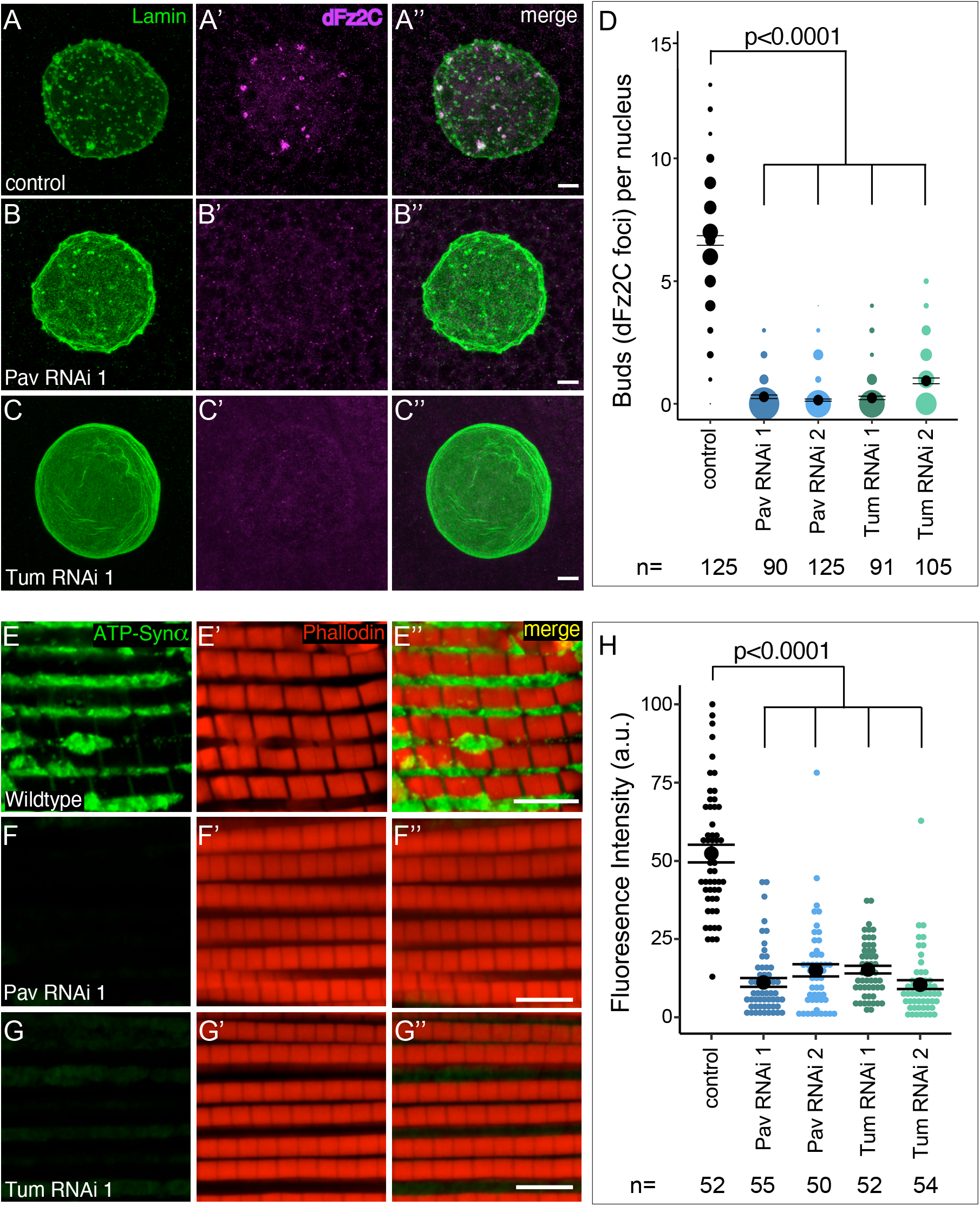
Pav and Tum knockdown nuclei lack NE buds and display NE bud associated phenotypes. (A-C”) Confocal micrograph projections of control (A-A”), Pav RNAi (B-B”), Tum RNAi (C-C”) larval salivary gland nuclei stained with Lamin B and dFz2C. (D) Quantification of NE buds per nucleus from two independent Pav and Tum RNAi lines. (E-G”) Confocal micrograph projection of adult indirect flight muscle (IFM) from control (+/Sgs-Gal4; E-E”), Pav RNAi (F-F”), Tum RNAi (G-G”) aged 21 days then stained with the activity dependent mitochondrial marker ATP-Syn α) and with Phalloidin. (H) Quantification of mitochondrial fluorescence intensity in IFM. Kruskal-Wallis test (D, H); all p-values indicated. Scale bars: 5μm.

Loss of NE-budding has been shown to result in aberrant NMJ development and loss of mitochondrial integrity (Jokhi et al., 2013; Li et al., 2016; Speese et al., 2012). We looked at whether Pav and Tum affect mitochondrial integrity in the indirect flight muscle (IFM) using the activity dependent mitochondrial marker ATP-Synthetase α. Consistent with their loss of dFz2C foci/NE buds, we find that both Pav and Tum knockdowns display the loss of mitochondrial integrity phenotype associated with NE budding (Fig. 2E-H; Fig. S2C-D”). Pav IFMs from adults aged 21 days show a 4.7 (n=55) and 3.6 (n=50) fold decrease, respectively, in mitochondrial activity compared to control (n=52; p<0.0001) (Fig. 2E-F”; H; Fig. S2C-C”). Similarly, Tum IFMs from adults aged 21 days show a 3.0 (n=54) and 4.5 (n=54) fold decrease, respectively, in mitochondrial activity compared to control (n=52; p<0.0001) (Fig. 2G-G”; H; Fig. S2D-D”). Taken together, these data indicate that Pav and Tum are essential players, and not cargos, of the NE budding process.

### Tum, but not Pav, functions in the Wash-SHRC nuclear complex

Consistent with its role in multiple nuclear processes, we find that *Drosophila* Wash is present in several nuclear complexes when we separated protein complexes from fly Kc cell nuclear lysates using blue native-PAGE (Fig. 3A). We showed previously that one major Wash-containing complex (~900KDa) includes the SHRC components, and is likely to be the Wash-containing complex required for its role in NE budding (cf. FAM21; Fig. 3A) (Verboon et al., 2020). Wash and the SHRC subunits only account for ~520kDa of the ~900kDa Wash/SHRC-containing complex, thus, we expected that Pav and Tum would be additional components of this complex. Surprisingly, only Tum overlaps with the Wash-SHRC complex (Fig. 3A). Interestingly, Tum also overlaps with a separate ~750kDa Wash and Pav-containing complex, which is the only nuclear Pav-containing complex (Fig. 3A).

**Fig. 3.**
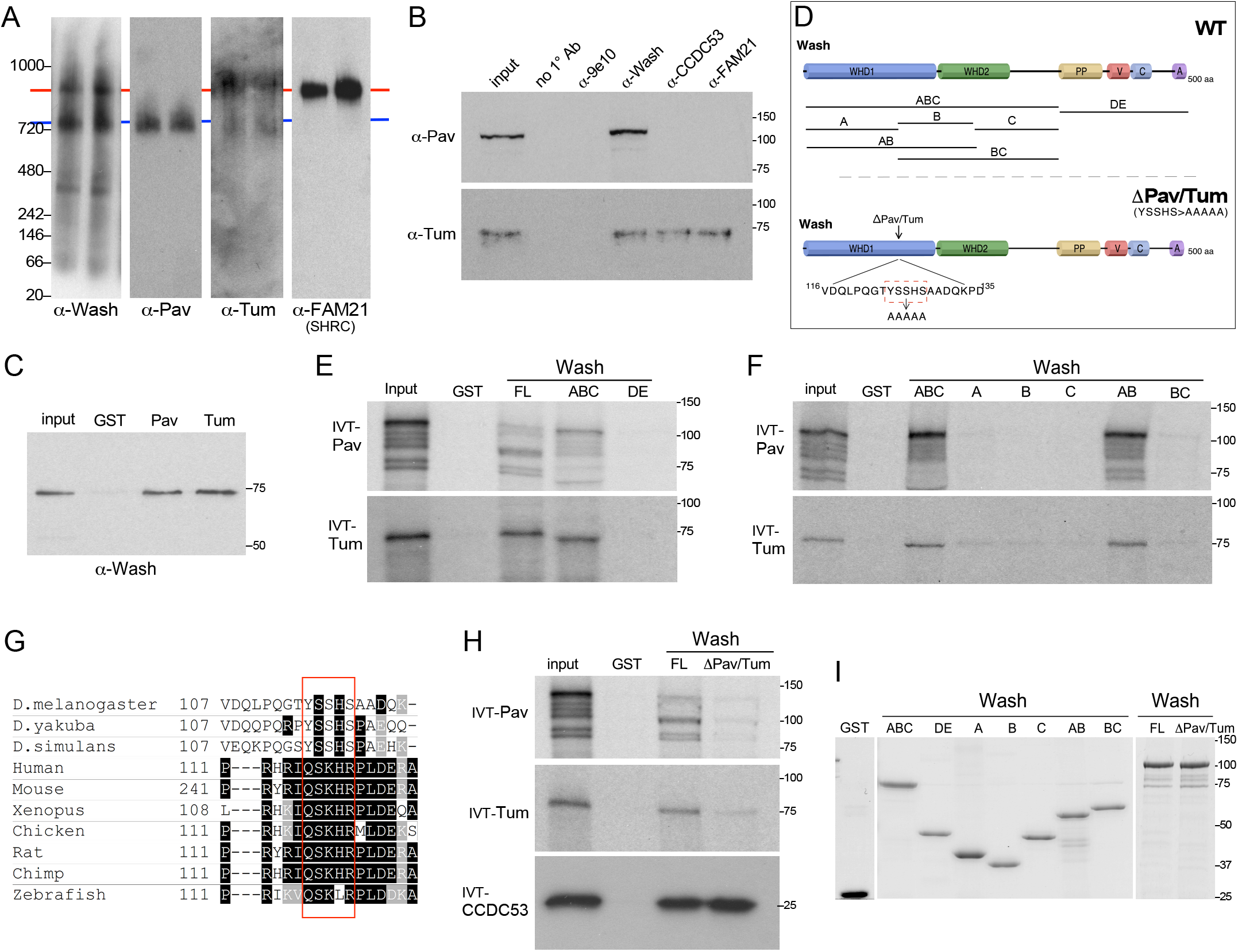
Pav and Tum interact with Wash in a conserved hydrophobic region within Wash’s WHD1 domain. (A) Western blots from Blue Native PAGE of Drosophila Kc cell nuclear extracts probed with antibodies to Wash, Pav, Tum, and FAM21 (SHRC subunit). Putative ~900KDa complex with Wash, Tum, and the SHRC (red line) and ~720KDa complex with Wash, Pav, and Tum (blue line) are indicated. (B) Western blots of immunoprecipitations from Kc cell nuclear extracts with no primary antibody included (no 1° Ab), a non-specific antibody (9e10), Wash, CCDC53, and FAM21. Blots were probed with antibodies to Pav and Tum as indicated. (C) GST pulldown experiments with bacterially purified proteins demonstrating Pav and Tum bind directly to full-length Wash. (D) Schematic diagram of the Wash wildtype (WT) rescue and substitution mutation constructs indicating the position and specific substitution mutations for the Wash^ΔPav/Tum^ construct and the Wash fragments used for mapping (not drawn to scale). (E-F) GST pulldown assays with ^35^S-labeled *in vitro* translated Pav and Tum and bacterially purified GST-Wash full length and fragments indicated. 10% input is shown. (G) Sequence alignment of the Wash region required for binding to Pav and Tum (red box). (H) GST pulldown assays with ^35^S-labeled *in vitro* translated Pav and Tum and bacterially purified GST-Wash^WT^ and GST-Wash^ΔPav/Tum^. 10% input shown. (I) Coomassie stained gel of purified GST fusion proteins used in this study to examine protein interactions among Wash, Pav, and Tum.

To confirm that Wash forms distinct nuclear complexes with Tum and Pav+Tum, we immunoprecipitated Wash and two SHRC subunits (CCDC53 and FAM21) from fly Kc cell nuclear lysates and probed the resulting western blots for Pav and Tum (Fig 3B). Consistent with the blue native-PAGE analyses, Pav is immunoprecipitated by Wash, but not by the SHRC components, whereas Tum is immunoprecipitated by Wash and the two SHRC components (Fig. 3B). Using bacterially purified proteins, we find that the interaction of Pav and Tum with Wash is direct (Fig. 3C). Thus, Wash appears to function in NE budding through at least two independent nuclear complexes.

### Pav and Tum bind to the same region of Wash

To further delineate Wash–SHRC–Tum and Wash–Pav–Tum complexes functions in NE budding, we mapped the sites on the Wash protein required for its binding to Pav and Tum using a series of Wash protein fragments (Fig. 3D). Unexpectedly, Pav and Tum mapped to the same region of the Wash protein: the junction of Wash pieces A and B (Fig. 3D-F, I), We then made a specific substitution mutation (YSSHS>AAAA) at this junction in the context of full-length Wash protein, designated Wash^ΔPav/Tum^. This Wash^ΔPav/Tum^ mutation reduced binding of both Pav and Tum to Wash in GST pulldown assays, whereas the SHRC component CCDC53 binds to Wash^ΔPav/Tum^ and Wash^WT^ (Fig. 3G-I; Fig. S2E).

### Wash–Pav–Tum interactions are required for NE-budding

To further delineate Wash–Pav–Tum functions in NE-budding we generated a transgenic line with the Wash^ΔPav/Tum^ substitution mutation under the control of the endogenous Wash promoter, then crossed it into the *wash* null homozygous background so that the only Wash activity comes from the transgene (*wash^ΔPav/Tum^*; see Methods). We have previously generated similar lines carrying a wildtype Wash transgene (*wash^WT^*), as well as ones disrupting Wash’s interaction with the SHRC component CCDC53 (*wash^ΔSHRC^*) and the Arp2/3 complex (*wash^ΔArp^*) (Verboon et al., 2020). Larval salivary gland nuclei from the from *wash^ΔPav/Tum^* mutants show an average of 0.02±0.0 dFz2C foci/NE-buds per nucleus (n=166) compared to 6.84±0.2 dFz2C foci/NE-buds per nucleus in the *wash^WT^* controls (n=122, p<0.0001) (Fig. 4A-C).

**Fig. 4.**
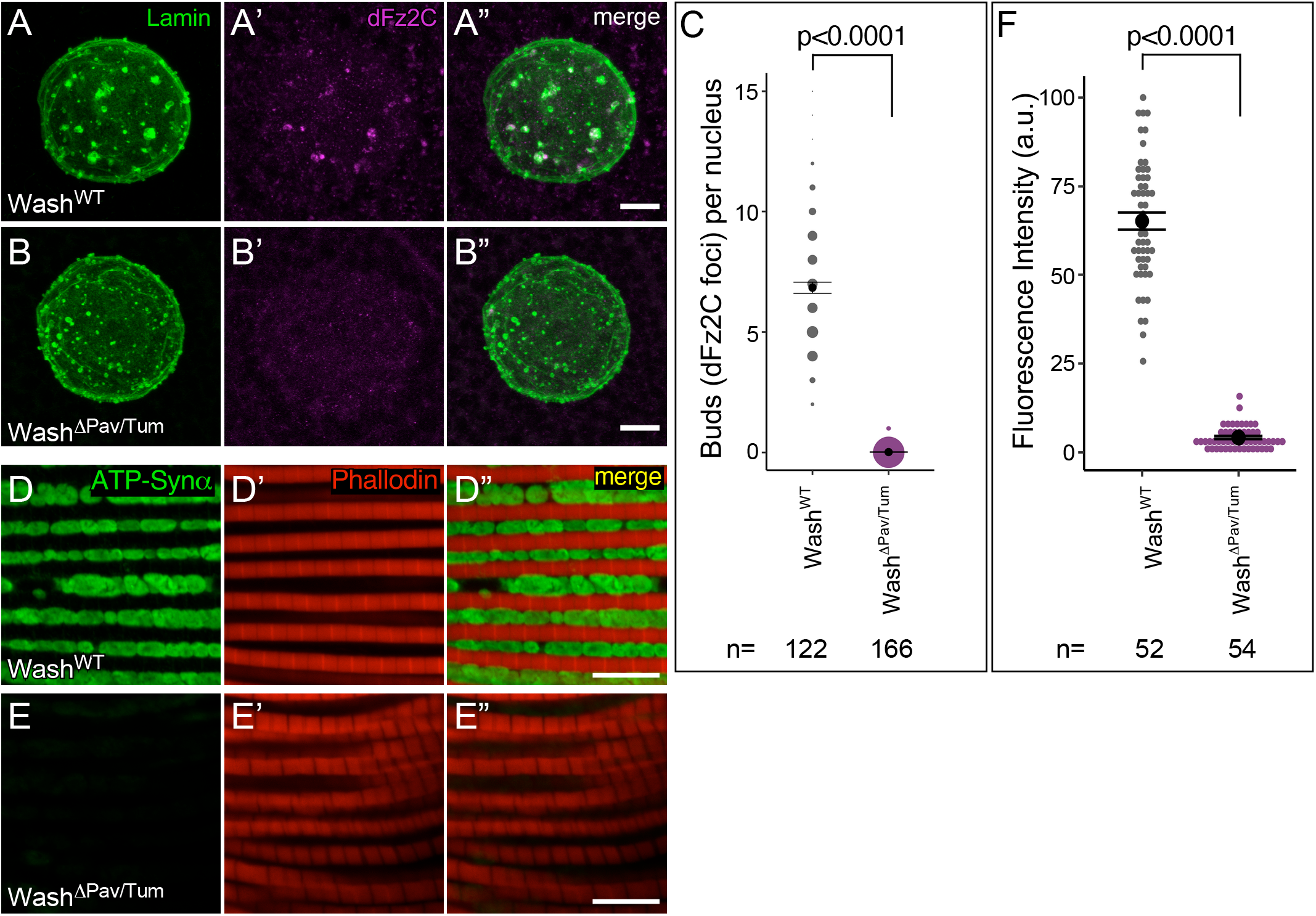
Wash binding to Pav and Tum is required for NE budding. (A-B”) Confocal micrograph projections of larval salivary gland nuclei from *wash*^WT^ (A-A”) and *wash*^ΔPav/Tum^ (B-B”) stained with Lamin B and dFz2C. (C) Quantification of NE buds per nucleus in larval salivary gland nuclei. (D-E”) Confocal micrograph projections of adult IFM from *wash*^WT^ (D-D”) and *wash*^ΔPav/Tum^ (E-E”) flies aged 21 days then stained for the activity dependent mitochondrial marker ATP-Syn α and with Phalloidin. (F) Quantification of ATP-Syn α fluorescence intensity from adult IFMs. Kruskal-Wallis test (C,F); all p-values indicated. Scale bars: 5μm.

As expected given its overall reduced number of NE buds, IFM from 21-day-old *wash^ΔPav/Tum^* show a decrease in mitochondrial activity, as assayed through assessing the level of ATP-Synthetase α (Fig. 4D-F). IFMs from *wash^ΔPav/Tum^* flies show a 15.8-fold decrease when compared to IFMs from the washWT construct (P<0.0001). Our data strongly suggest that Wash works together with Pav and Tum in NE budding.

### Wash requires its bundling activity to form NE buds

WAS family proteins have been implicated in membrane deformations (i.e. protrusions and endocytosis) in the cytoplasm given its branched actin nucleation activity and its interactions with the cortical cytoskeleton and overlying plasma membrane (Burianek and Soderling, 2013; Campellone and Welch, 2010; Massaad et al., 2013; Rottner et al., 2010; Rotty et al., 2013; Stradal et al., 2004; Takenawa and Suetsugu, 2007). We showed previously that Wash’s branched actin nucleation activity is required for NE bud formation through its ~900 KDa Wash–SHRC-Tum complex (Verboon et al., 2020). We have also shown that Wash encodes several context-dependent biochemical activities in addition to its branched actin nucleation functions, including actin and microtubule (MT) binding, bundling, and crosslinking (Liu et al., 2009; Verboon et al., 2018; Verboon et al., 2020; Verboon et al., 2015a; Verboon et al., 2015b). Since the ~750 KDa Wash-Pav-Tum complex does not appear to be associated with branched actin nucleation activity, we examined whether Wash’s actin bundling activity is required for NE bud formation using *in vitro* F-actin bundling assays.

*In vitro* polymerized F-actin distributes uniformly in the absence of additional protein (Fig. 5A) Addition of bacterially purified full-length Wash^WT^ protein bundles actin, as previously reported (Fig. 5B) (Liu et al., 2009). Similarly, addition of Wash^ΔSHRC^ or Wash^ΔArp2/3^ proteins bundles actin (Fig. 5C-D). Suprisingly, addition of Wash^ΔPav/Tum^ does not bundle actin (Fig 6E). The inability of Wash^ΔPav/Tum^ to bundle actin is not due to its inability to bind F-actin. We find that both bacterially purified GFP-Wash^WT^ and GFP-Wash^ΔPav/Tum^ can bind to pre-bundled F-actin (bundled using the formin homology protein Cappuccino; (Rosales-Nieves et al., 2006)) (Fig. 5F-G”). Thus, in addition to Wash’s branched actin nucleation activity, these results suggest that Wash may require its F-actin bundling activity during NE budding.

**Fig. 5.**
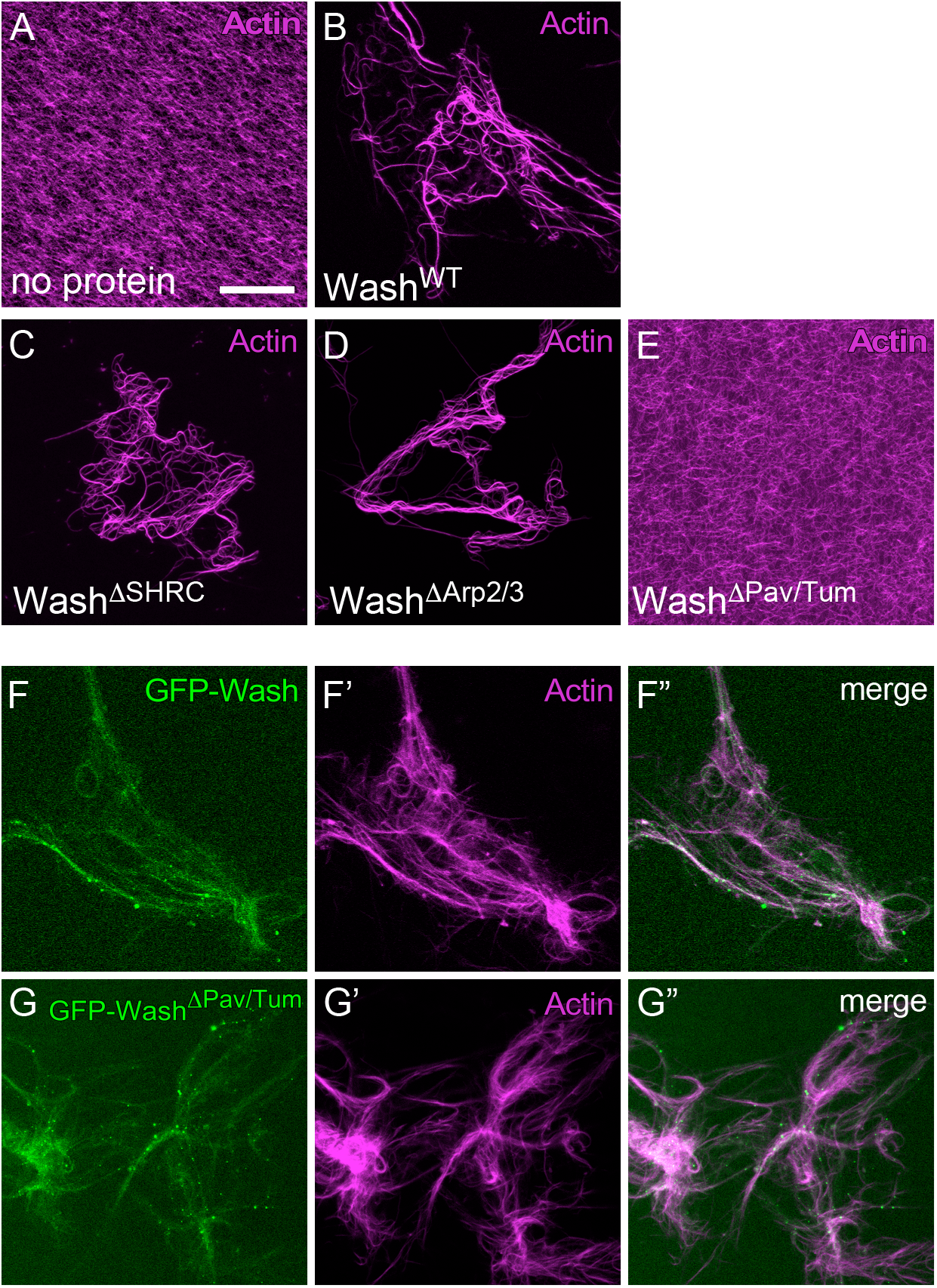
The *wash*^ΔPav/Tum^ mutation binds, but can no longer bundle, actin. (A-E) Phalloidin-stabilized actin filaments incubated with no protein (A), Wash^WT^ (B), Wash^ΔSHRC^ (C), Wash^ΔArp2/3^ (D), or Wash^ΔPav/Tum^ (E) proteins. Note the lack of actin bundling in the presence of Wash^ΔPav/Tum^ protein. (F-G”) Binding of sfGFP-Wash (F-F”) and sfGFP-Wash^ΔPav/Tum^ (G-G”) to phalloidin-stabilized actin filaments that were bundled with Capu (formin) protein. Final protein concentrations for bundling assays: Wash^WT^, 500 nM; Wash^ΔSHRC^, 500 nM; Wash^ΔArp2/3^, 500 nM; Wash^ΔPav/Tum^, 500 nM; CapuFH2, 500 nM; sfGFP-Wash, 50 nM; and sfGFP-Wash^ΔPav/Tum^, 50 nM. Scale bar: 30 μm

**Fig. 6.**
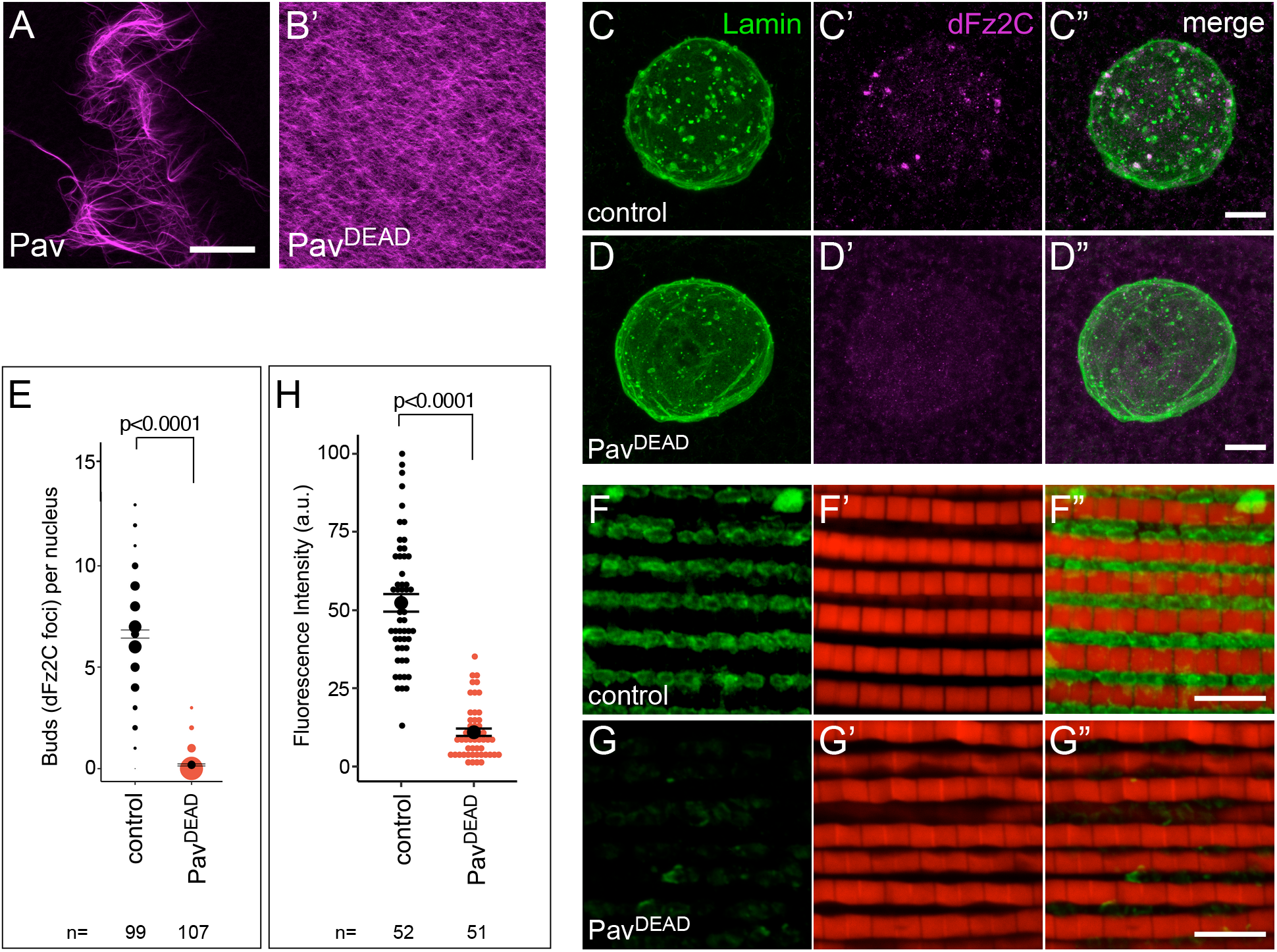
Bundled actin is required for NE bud formation. (A-B) Phalloidin-stabilized actin filaments were incubated with Pav^WT^ (A) and Pav^DEAD^ (B) protein. Final protein concentrations for bundling assays: Pav^WT^, 200 nM; Pav^DEAD^, 200 nM. (C-D”) Confocal micrograph projections of larval salivary gland nuclei from control (C-C”) and Pav^DEAD^ (D-D”) stained with Lamin B and dFz2C. (E) Quantification of NE buds per nucleus in larval salivary gland nuclei. (F-G”) Confocal micrograph projections of adult IFM from control (F-F”) and Pav^DEAD^ (G-G”) flies aged 21 days then stained for the activity dependent mitochondrial marker ATP-Syn α and Phalloidin. (H) Quantification of ATP-Syn α fluorescence intensity from adult IFMs. Kruskal-Wallis test (E, H); all p-values indicated. Scale bars: 30 μm in A-B; 5μm in C-G”.

### Pav requires its non-canonical F-actin bundling activity during NE budding

We previously identified a non-canonical role of the kinesin-like protein, Pav, in wound healing and oogenesis: in addition to its MT binding activity, Pav also binds and bundles F-actin (Nakamura et al., 2020). Interestingly, we found that a G131E mutation in Pav (referred to as Pav^DEAD^; (Minestrini et al., 2002)) that exhibits rigor-like association to and stabilization of MTs *in vivo*, can bind to but no longer bundle F-actin (Fig. 6A-B) (Nakamura et al., 2020). We hypothesized that if F-actin bundling is required for NE bud formation, then Pav^DEAD^, like Wash^ΔPav/Tum^, should exhibit NE bud phenotypes. Larval salivary gland nuclei from the Pav^DEAD^ mutant shows an average of 0.02±0.52 dFz2C foci/NE-buds per nucleus (n=107) compared to 6.64±2.1 dFz2C foci/NE-buds per nucleus in the control (n=99, p<0.0001) (Fig. 6C-E). As expected with its overall reduction in NE buds, IFM from 21-day-old Pav^DEAD^ adults show a 4.8-fold decrease in mitochondrial activity compared to control adults (p<0.0001) (Fig. 6F-H). These data strongly suggest Wash’s F-actin bundling activity requires its interaction with Pav and that this F-actin bundling activity is required in the NE budding process.

To further delineate the role of actin nucleation and bundling in the process of NE budding, we treated larval salivary glands co-expressing GFP-Lamin and dFz2-Scarlet with Latrunculin B (Lat B; inhibitor of actin polymerization) (Fig. 7A-E). LatB inhibition of actin polymerization immediately (< 30 sec) disrupts dFz2C localization to NE-buds (Fig. 7D-E) compared to DMSO control (Fig. 7C, E), however consistent with Wash-Lamin indirect effects on NE budding, lamin localization to the inner NE-bud periphery remains intact (Fig. 7A-B, E). We are unable to examine the effect of LatB on Pav or Tum expression in NE-buds due to masking by their unrelated nuclear/Centralspindlin roles (data not shown). Altogether, our data shows both actin polymerization and bundling are required for formation of NE buds, and in particular, F-actin bundling activity is likely required for the organization of macromolecules within NE buds.

**Fig. 7.**
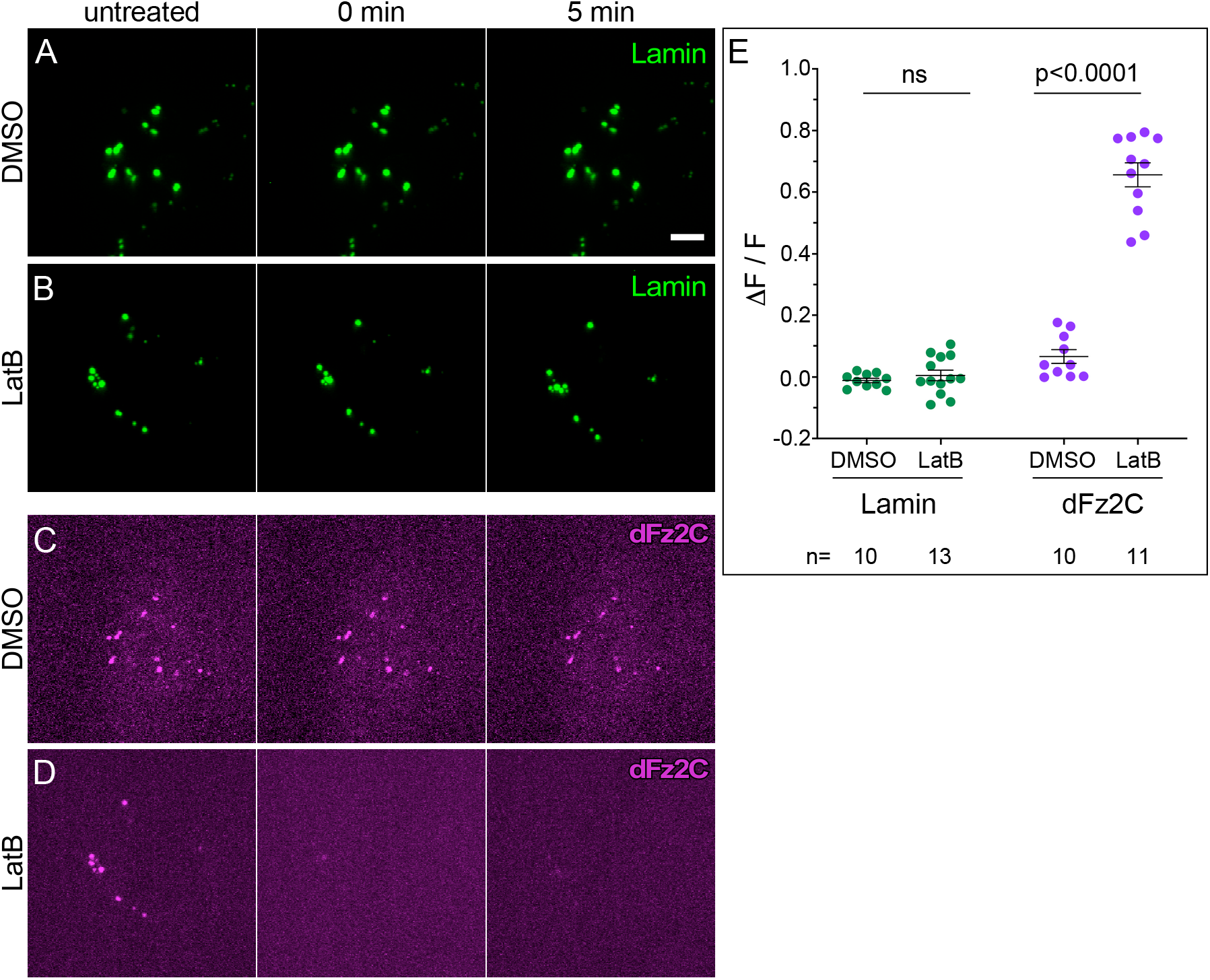
Retention of dFz2C within NE buds requires actin. Time-lapse confocal xy projection images from *Drosophila* salivary gland nuclei expressing GFP-Lamin (A-B) or dFz2-Scarlet (C-D) treated with DMSO (A, C) or Latrunculin B (LatB) (B, D) at the times indicated. (E) Quantification of the GFP-Lamin and dFz2-Scarlet in the conditions indicated. Two-tailed student’s t-test; all p-values indicated. Scale bars: 10μm.

## DISCUSSION

NE-budding is an evolutionarily conserved process that provides an alternate, nuclear pore-independent, route out of the nucleus for large macromolecules, using a nuclear envelope budding mechanism reminiscent of HSV-1 nuclear egress. We previously showed that Wash, its regulatory complex (SHRC), Arp2/3, and actin capping protein are required for the formation of NE buds in *Drosophila* salivary gland nuclei. Here, we identify Pav (MKLP1/Kif23) and Tum (RacGAP50C/RacGAP1) as new components of Wash-containing complexes involved in NE bud formation and organization. Tum, but not Pav, is part of the ~900 KDa Wash-containing nuclear complex previously linked to NE budding (Verboon 2020). Wash, however, forms a second ~750KDa NE budding-associated nuclear complex that contains both Pav and Tum. Interestingly, Pav and Tum bind to the same site in the Wash protein. We also find that Wash’s actin nucleation and bundling activities are required for distinct roles in the formation of NE buds and organization/maintenance of macromolecules within NE buds. Thus, Wash is present in several independent nuclear complexes. We have examined three of these in NE budding and find: 1) A ~450KDa Wash-LaminB complex <u>indirectly</u> regulates NE bud formation through its general disruption of the nuclear lamina leading to inefficient NE bud formation (Verboon et al., 2020). 2) A ~900KDa Wash-SHRC-Tum complex interacts with Arp2/3 to <u>nucleate</u> branched actin filaments required for NE bud formation. 3) A ~750KDa Wash-Pav-Tum complex provides infrastructure to NE buds via its actin <u>bundling</u> regulation. Thus, Pav and Tum are providing exciting new entry points into the physical machineries of this relatively new nuclear export pathway.

### The roles of actin nucleation and bundling during the NE-budding process

The cortical actin cytoskeleton has been implicated in scaffolding exocytic/endocytic deformations by underlying the plasma membrane such that it can bend membranes, anchor cargos within the deformations, and/or provide internal organization/infrastructure to the vesicles/protrusions formed (Blanchoin et al., 2014; Buracco et al., 2019; Chakrabarti et al., 2021; Gundelfinger et al., 2003; Kessels and Qualmann, 2021). Branched actin nucleation by WAS proteins and the Arp2/3 complex near the membrane can generate the force necessary to deform membrane and the underlying cytoskeleton in many cellular processes, including endocytosis and the formation of cellular protrusions, whereas actin bundling acts to assemble and organize/scaffold proteins within a cell (Rotty et al., 2013; Siton-Mendelson and Bernheim-Groswasser, 2017; Svitkina, 2018; Swaney and Li, 2016). Interestingly, the encapsulation or confinement of RNAs or proteins within vesicles has been shown to promote RNA-RNA and RNA-protein associations, organization, stability, and functions (Cheng et al., 2018; Peng et al., 2021; Ross et al., 2020). Our previous study suggests that actin organization is indispensable for the formation of NE buds. While we expected that LatB treatment would disrupt all NE bud structures, actin disruption affected dFz2 foci, but not Lamin foci. We showed the inhibition of actin dynamics with LatB treatment leads to the immediate loss of dFz2C foci from NE buds, suggesting actin scaffolding is required for the organization of cargos and/or machineries within NE buds. Thus, Pav and Tum may work within the NE bud to aide in the final organization and/or maturation of megaRNP granules prior to their release into the cytoplasm.

Notably, Lamin foci remained intact after LatB treatment, further confirming the indirect and independent role of the Wash-Lamin complex on NE bud formation (Verboon et al., 2020), and strengthening our conclusions that Wash’s nucleation and/or bundling activities likely scaffold NE-buds allowing for the proper recruitment of cargo and machinery. Our findings suggest that the two independent Wash-containing complexes required for the NE-budding process have distinct roles (Fig. 8): 1) the ~900 KDa Wash-SHRC-Tum complex works with Arp2/3 to nucleate branched actin filaments that are required for the nuclear envelope deformations needed for NE bud formation and the inclusion of cargos, and 2) the ~750KDa Wash-Pav-Tum complex establishes a scaffold-like organization within NE buds that stabilizes, organizes, and/or anchors megaRNP complexes or other large macromolecular complexes through the regulation of actin bundling. This is consistent with our previous electron microscopy studies where we observed a subset of NE buds form prior to cargo entry (Verboon et al., 2020). In addition, Pav can bundle actin, whereas Tum cannot (Nakamura et al., 2020). While we find that disruption of Pav’s bundling activity by overexpressing Pav^DEAD^ impairs the formation of NE buds, the formation of megaRNPs at the bud site might be required for the initiation of NE bud formation. Developing an inducible live imaging model combined with genetic and pharmacological approaches will be needed to understand the precise spatial and temporal regulation of actin dynamics, nuclear envelope deformation, and megaRNP formation during NE-budding process.

**Fig. 8.**
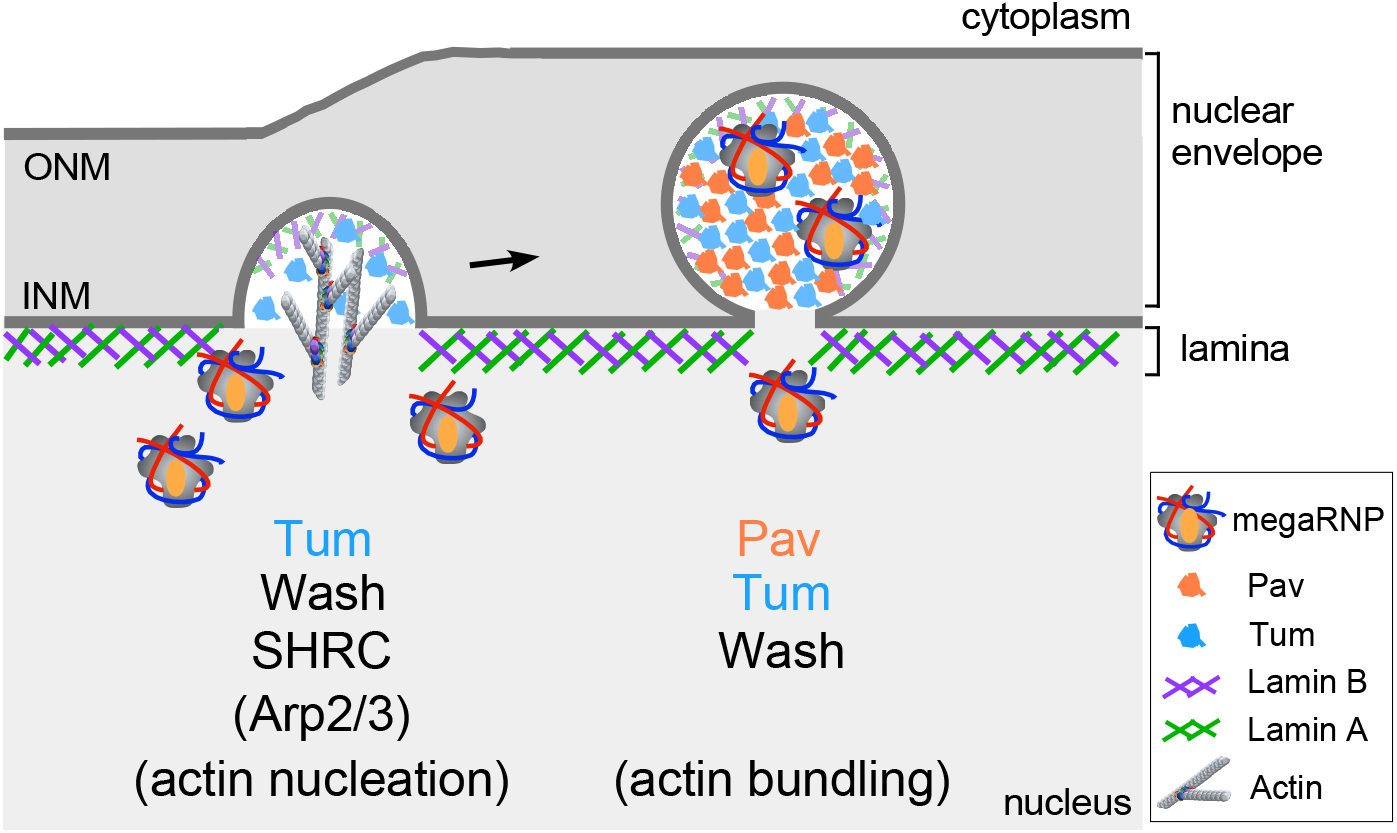
Roles of Wash, Pav, and Tum in NE bud formation and/or organization. Schematic diagram of NE-budding highlighting the proposed roles of the ~900 KDa Wash/SHRC/Tum complex in NE-bud formation and the ~750 KDa Wash/Pav/Tum complex in NE-bud internal infrastructure/organization.

### Functions of Pav, Tum, and Wash in NE budding and the regulation of Wnt signaling

The NE budding pathway was revealed as an alternate nuclear export route during pioneering studies investigating Wnt-dependent neuro-muscular junction synapse development in *Drosophila* body wall muscles (Speese et al., 2012). In this context, *Drosophila* Wnt-1, Wingless (Wg), secreted from motor neuron cells activates the dFrizzled-2 (Wg receptor) in muscle cells. Following internalization, a C-terminal portion of the receptor protein is cleaved (dFz2C) and transported into the nucleus where it is incorporated into megaRNPs that exit the nucleus by budding through the nuclear envelope (Speese et al., 2012). Consistent with this, *wash* mutants exhibit a dominant zygotic synthetic lethality when trans-heterozygous with *wg* mutants (SMP, unpublished observation). Interestingly, Pav and Tum have also been shown to have a non-canonical nuclear role in Wg/Wnt regulation, where they negatively regulate canonical Wg/Wnt signaling downstream of β-catenin stabilization, where they are proposed to antagonize Wnt target gene expression (Jones et al., 2010). This study also showed that some of Tum’s regulatory function is Pav-independent: Tum protein that cannot bind to Pav (TumΔPav) retained a modest ability to repress Wg/Wnt signaling in cultured cells (Jones et al., 2010). This is consistent with our finding that Tum forms Pav-dependent and Pav-independent Wash-containing complexes in the nucleus.

While the transcriptional antagonism observed with Pav or Tum may be due to the inability to form active transcriptional complexes at Wg/Wnt target gene promoters (Jones et al., 2010), given the roles of Pav and Tum in NE budding, it may also result from the inability of particular protein complexes to reach their target sites. Since dFz2C foci/NE buds are not formed in the absence of Pav, Tum, Wash, or Wash^ΔPav/Tum^, this would likely alter the number, types, and/or activities of Pav, Tum, and/or Wash complexes formed in the nucleus resulting in a functional imbalance.

Such a disruption of functional complexes would be important not only for normal developmental events, but for cancers and virus infections as well. Over-expression of Kif23 (Pav) activates canonical Wnt signaling in gastric and colorectal cancer cells, whereas Kif23 knockdown suppresses cell proliferation, tumorigenesis, migration, and invasion (Ji et al., 2021; Liu et al., 2020). Tum (RACGAP1) is upregulated in prostate, ovarian, and breast cancers (Ren et al., 2021; Song et al., 2019; Wang et al., 2018). *Drosophila* Pav and Tum have also been shown to recruit the Alix and ESCRT III proteins to the midbody during cytokinesis using a mechanism analogous to that involved in retroviral budding through the plasma membrane (Lie-Jensen et al., 2019), although a recent study of NE budding in yeast showed that ESCRTIII proteins did not appear to contribute to NE budding events (Panagaki et al., 2021). NE budding also shares many similarities with herpesvirus nuclear egress mechanisms (Bigalke and Heldwein, 2015; Bigalke and Heldwein, 2016; Fradkin and Budnik, 2016; Johnson and Baines, 2011; Lee and Chen, 2010; Mettenleiter et al., 2013; Parchure et al., 2017). Interestingly, recent studies have shown that HSV-1 infection is productively stimulated when VP16 enhances canonical Wnt signaling activity (Zhu and Jones, 2018), and Wnt antagonists prevent HSV-1 replication (Harrison and Jones, 2021). Thus, NE budding may be a common regulatory theme and a new area of exploration for future HSV-1 infection therapies.

Thus, nuclear export via NE budding is an important new finding with widespread implications for regulating fundamental cellular/developmental processes and aging, as well as diseases such as neuropathies, cancer progression, and potentially cell-virus interactions. These studies provide exciting new avenues in Wash, Pav, and Tum for investigating the structural components of NE-budding and for understanding general aspects of nucleo-cytoplasmic transport, as well as for exploring the developmental events regulated by this process, cancer proliferation, and HSV-1 virus budding mechanisms.

## MATERIALS AND METHODS

### Reagents/Resources

Specific information for the reagents and resources used in this study are given in Table S1.

### Fly stocks and genetics

Flies were cultured and crossed at 25°C on yeast-cornmeal-molasses-malt extract medium. Flies used in this study are listed in Table S1. All fly stocks were treated with tetracycline and then tested by PCR to ensure that they did not harbor Wolbachia. RNAi knockdowns were driven in the salivary glands by the GAL4-UAS system using the P{Sgs3-GAL4.PD} driver (Bloomington Drosophila Stock Center, stock #6870). RNAi knockdowns were driven in the indirect flight muscle by the GAL4-UAS system using the P{w[+mC]=Mhc-GAL4.K2} driver (Bloomington Drosophila Stock Center, stock #55133). The *wash^Δ185^* deletion allele was kept as a continuously outcrossed stock (Verboon et al., 2018).

### Construction of Wash^ΔPav/Tum^ mutant transgenic lines

The residues required for the interaction of Wash with Pav and Tum were mapped by successive GST pulldown assays using fragments of Wash protein, followed by the generation of a specific substitution mutation (YSSHS>AAAAA) in the context of the full-length Wash protein (as detailed in Fig. 4D-H). This Wash^ΔPav/Tum^ substitution mutation was confirmed by sequencing. As previously described, a 2.9-kb genomic fragment encompassing the entire *wash* gene was amplified by PCR, then subcloned into the Casper 4 transformation vector by adding KpnI (5’) and BamHI (3’) restriction sites. GFP was inserted N-terminal to the Wash ATG by PCR (GFP-Wash^WT^) (Verboon 2018). The Wash portion of this construct was swapped with the substitution mutations described above to generate GFP-Wash^ΔPav/Tum^. The Wash^ΔPav/Tum^ substitution mutation was confirmed by sequencing.

The GFP-Wash^ΔPav/Tum^ construct was used to make germline transformants as previously described (Spradling, 1986). Transgenic lines that mapped to chromosome 2 and that had non-lethal insertions were kept. The resulting transgenic lines P{*w^+^; GFP-Wash^ΔPav/Tum^*) were recombined onto the *wash^Δ185^* null chromosome to assess the contribution of the mutant Wash transgene. The resulting recombinants (*wash^Δ185^* P{*w^+^*; *GFP-Wash^WT^}* and *wash^Δ185^* P{*w^+^*; *GFP-Wash^ΔPav/Tum^}*) are essentially gene replacements, as Wash activity is only provided by the transgene. These transgenes do not rely on overexpression, but rather on the spatial and temporal expression driven by the endogenous *wash* promoter itself. We analyzed a minimum of three independent transgenic lines per construct and checked all lines to confirm that the levels and spatial distribution of their expression is indistinguishable from that in wildtype. The wildtype version of this transgene (*wash^Δ185^* P{*w^+^*; *GFP-Wash^WT^}*) rescues the phenotypes associated with the outcrossed *wash^Δ185^* mutation (Liu et al., 2009; Verboon et al., 2018; Verboon et al., 2020; Verboon et al., 2015a; Verboon et al., 2015b).

### Lysate preparation

*Drosophila* cytoplasmic and nuclear extracts were made from Kc167 cells. Briefly, cells were grown to confluence in 500 ml spinflasks, pelleted for 5 min at 500 g, resuspended in 100 ml cold 1× PBS and re-pelleted for 5 min at 500 g. Cell pellets were flash frozen in liquid nitrogen. Cells were resuspended in sucrose buffer (0.32 M sucrose, 3 mM CaCl_2_, 2 mM MgAc, 0.1 mM EDTA, 1.5% NP40) with 2× protease inhibitors [Complete protease inhibitor (EDTA free; Sigma, St Louis, MO), 2 mM PMSF and 1 mM Na_3_VO_4_] and 2× phosphatase inhibitors [PhosSTOP; Sigma, St Louis, MO] at 100 μl per 1 million cells and incubated on ice for 30 min. Lysate was dounce homogenized 10× on ice. Lysate was then centrifuged 10 min at 2900 g at 4°C, nuclei formed a pellet and supernatant was cytoplasmic extract. Lipids were removed from the top of cytoplasmic extract using a sterile swab, then the cytoplasmic fraction was removed and centrifuged for 10 min at 3300 g at 4°C. Cytoplasmic supernatant was removed and one-tenth of the supernatant volume of 11× RIPA was added. Cytoplasmic extract was aliquoted and flash frozen. Nuclear pellet was resuspended with sucrose buffer with protease/phosphatase inhibitors and NP40 and re-dounced. Nuclear lysate was then centrifuged for 10 min at 3300 g at 4°C, and supernatant was discarded. The nuclear pellet was resuspended in sucrose buffer without NP40 and centrifuged for 20 min at 3300 g at 4°C. The supernatant was discarded and nuclear pellet was resuspended in 2.5 ml of buffer (20 mM HEPES pH 7.9, 0.5 mM EDTA, 100 mM KCl and 10% glycerol) per liter of cells used. DNA was degraded by incubating with MNase at 37°C for 10 min. 20 μl of 500 mM EDTA per 500 μl lysate was added and incubated on ice for 5 min. Lysate was then nutated for 2 h at 4°C. Lysate was sonicated using a Sonic Dismembrator (Model 60; Fisher Scientific) at setting 3.5 with 10 s per pulse for 15 min. Lysate was clarified with a 15 min centrifugation at 25,000 g, aliquoted and flash frozen. Cytoplasmic and nuclear purity of lysates was assayed by western blot as previously described (Verboon et al., 2020). Donkey anti-mouse or anti-rabbit HRP (1:15,000, Jackson ImmunoResearch Labs) secondary antibodies were used.

### Pav antibody production and characterization

Due to the limited availability of Pav antibody (Adams et al., 1998), we generated mouse polyclonal and monoclonal antibodies to Pav. For expression of Pav protein, a DNA fragment covering amino acids 208–687 of Pav was generated by PCR and cloned into a pET21 vector. Protein was induced as described previously (Rosales-Nieves et al., 2006). Induced protein was purified with Fastflow Nickel-Sepharose (GE) under denaturing conditions. Purified Pav protein was dialyzed into PBS. Pav polyclonal sera and monoclonal lines (L24 and O8) were generated in the Fred Hutch Antibody Technology Shared Resource Facility. Western blotting was used to test monoclonal antibody specificity against *in vitro* translated full-length Pav protein, wildtype embryo whole cell extract, Kc cell nuclear extract, and Kc cell cyto extract. *Drosophila* whole cell (from 0- to 2-hour embryos) extract was a gift from Toshi Tsukiyama (Fred Hutch). Secondary antibody used was donkey anti-mouse HRP (1:15,000 dilution; Jackson ImmunoResearch).

### Immunostaining of embryos

0-12 hour embryos were fixed with 4% formaldehyde/heptane for 30 min and devitellinized by heptane/methanol. Immunofluorescence was performed as described previously (Abreu-Blanco et al., 2011). Mouse monoclonal L24 or O8 anti-Pav (1:10) were used as primary antibodies. Goat anti-mouse alexafluor-568 was used as secondary antibody. Embryos were mounted in SlowFade Gold (Invitrogen).

### Immunoprecipitation

Nuclear lysate was incubated with primary antibody overnight at 4°C. Protein G–Sepharose (20 μl) was then added in 0.5 ml Carol buffer (50 mM HEPES pH 7.9, 250 mM NaCl, 10 mM EDTA, 1 mM DTT, 10% glycerol, 0.1% Triton X-100) plus 0.5 mg/ml bovine serum albumin (BSA) and protease inhibitors (Complete EDTA-free Protease Inhibitor cocktail; Sigma) and the reaction allowed to proceed for 2 h at 4°C. The beads were washed 1× with Carol buffer plus BSA and 2× with Carol buffer alone. Analysis was conducted using SDS-PAGE followed by western blotting. Antibodies used for immunoprecipitations are as follows: anti-9e10 (1:9; Developmental Studies Hybridoma Bank), anti-Wash monoclonal (1:6; P3H3; (Rodriguez-Mesa et al., 2012)), anti-CCDC53 (1:1000; (Verboon et al., 2015a)), anti-Lamin B monoclonal (1:8; AD67.10, Developmental Studies Hybridoma Bank). Antibodies used for the IP western blots are as follows: rabbit anti-Tumbleweed polyclonal (1:2500; (Tao et al., 2016)), rabbit anti-Pavarotti (1:5000; (Adams et al., 1998)), and donkey anti-rabbit HRP (1:30,000, Jackson ImmunoResearch Labs).

### GST pulldown assays and Blue Native PAGE

GST pulldown assays were performed as previously described (Magie and Parkhurst, 2005; Magie et al., 2002; Rosales-Nieves et al., 2006).

Blue Native Page was performed using a Novex Native PAGE Bis-Tris Gel System (Invitrogen) following manufacturer protocols. Briefly, *Drosophila* Kc cell nuclear extract was centrifuged at 16,200 g for 15 min at 4°C, then 4× NativePAGE Sample Buffer (Invitrogen) was added to the supernatant. These prepared samples were loaded onto 3–12% Bis-Tris Native PAGE gels and electrophoresed using a 1× native PAGE running buffer system (Invitrogen). The cathode buffer included 1× cathode buffer additive (Invitrogen). Native mark protein standard (Invitrogen) was used as the molecular mass marker. The following antibodies were used: mouse anti-Wash monoclonal (P3H3, 1:2; (Rodriguez-Mesa et al., 2012)), mouse anti-Fam21 polyclonal, (1:400; (Verboon et al., 2015a)), rabbit anti-Tumbleweed (1:2500; (Tao et al., 2016)), rabbit anti- Pav (1:3000, (Adams et al., 1998)), and donkey anti-mouse or anti-rabbit HRP (1:30,000, Jackson ImmunoResearch Labs).

### Immunostaining of larval salivary glands

Salivary glands were dissected, fixed, stained and mounted as previously described (Verboon et al., 2015a). Primary antibodies were added at the following concentrations: mouse anti-Lamin B monoclonal (AD67.10; 1:200; Developmental Studies Hybridoma Bank), guinea pig anti-Fz2C (1:2500; (Verboon et al., 2020)), rabbit anti-Pav (1:250; (Adams et al., 1998)); mouse anti-Pav (polyclonal; 1:250; this study); mo use anti-Pav (monoclonal L24; 1:10; this study); rabbit anti-Tumbleweed (1:2500; (Tao et al., 2016)), rabbit anti-GFP (1:1000; Invitrogen).

### Live imaging of salivary gland nuclei

To generate the UASp-dFz2-Scarlet construct, the full-length dFz2 ORF was amplified and then mScarlet-I was fused to its 3’ end (Addgene #85044). The dFz2-Scarlet fusiomn was then cloned into the UASp vector. This UASp-dFz2-Scarlet construct was used to make germline transformants as previously described (Spradling, 1986).

Salivary glands were dissected in Schneider’s Drosophila medium (Gibco) and then placed in a glass bottom dish. For drug treatment, DMSO (control) or Latrunculin B (10 μM at final concentration; EMD Millipore) were added into medium. Images were acquired with i) a Revolution WD system or ii) an Ultraview Vox spinning disk confocal system. The Revolution WD system (Andor Technology Ltd., Concord, MA) is mounted on a Leica DMi8 (Leica Microsystems Inc., Buffalo Grove, IL) with a 63x/1.4 NA objective lens and controlled by MetaMorph software. Images were acquired using a 488 nm, 561 nm, and 633 nm Lasers and Andor iXon Ultra 888 EMCCD camera (Andor Technology Ltd.).

To compare the intensity of Lamin-GFP and dFz2C-Scarlet following drug treatment, we measured the mean intensity in the NE-buds before and 5 minutes after adding DMSO or LatB. ΔF/F is an averaged intensity change from each bud in the nucleus and each intensity change was calculated by (mean intensity before drug treatment - mean intensity after drug treatment) / mean intensity before drug treatment).

### Immunostaining of indirect flight muscle

UAS controlled RNAi-expressing flies were crossed to MHC-GAL4 driver flies and female RNAi/MHC-GAL4 trans-heterozygous flies were collected and aged 21 days at 25°C. Fly thoraxes were dissected and cut along the ventral side in cold PBS. Thoraxes were fixed using 1:6 fixative and heptane for 15 min. The fixative used was: 16.7 mM KPO_4_ pH 6.8, 75 mM KCl, 25 mM NaCl, 3.3 mM MgCl_2_ and 6% formaldehyde. After three washes with PTW (1× PBS, 0.1% Tween-20) thoraxes were then cut along the dorsal side resulting in two halves and fixed again for 10 min using 1:6 fix/heptane for 15 min. After three washes with PTW, thoraxes were permeabilized in 1× PBS plus 1% Triton X-100 for 2 h at room temperature, then blocked using PAT (1× PBS, 0.1% Tween- 20, 1% BSA, 0.05% azide) for 2 h at 4°C. The thoraxes were incubated in mouse anti-ATP-Synthase α (15H4C4; 1:100; Abcam, Cambridge, UK) for 48 h at 4°C. Thoraxes were washed three times with XNS (1× PBS, 0.1% Tween-20, 0.1% BSA, 2% normal goat serum) for 30 min each. Alexa Fluor-conjugated secondary antibodies (Invitrogen, Carlsbad, CA) diluted 1:1000 in PbT (1× PBS, 0.1% Tween-20, 0.1% BSA)) and Alexa Fluor-conjugated phalloidin (1:50) were then added and the thoraxes were incubated overnight at 4°C. Thoraxes were washed ten times with PTW at room temperature for 10 min each and were mounted on slides in SlowFade Gold medium (Invitrogen, Carlsbad, CA) and visualized using a Zeiss confocal microscope as described below. To quantify ATP-Synthase α expression, we measured 512×512 pixel regions of the IFM and measured total fluorescence using ImageJ (NIH). Fluorescence measurements from separate experiments were normalized to the control genotype.

### Microscopy

Images of fixed tissues were acquired using a Zeiss LSM 780 spectral confocal microscope (Carl Zeiss Microscopy GmbH, Jena, Germany) fitted with a Zeiss 40×/1.0 oil Plan-Apochromat objective and a Zeiss 63×/1.4 oil Plan-Apochromat objective. FITC (Alexa Fluor 488) fluorescence was excited with the 488 nm line of an argon laser, and detection was between 498 and 560 nm. Red (Alexa Fluor 568) fluorescence was excited with the 561 nm line of a DPSS laser and detection was between 570 and 670 nm. The pinhole was set to 1.0 Airy Units. Confocal sections were acquired at 0.2–1.0 μm spacing. Super-resolution images were acquired using an Airyscan detector in Super Resolution mode and captured confocal images were then processed using the Airyscan Processing feature on the Zen software provided by the manufacturer (Carl Zeiss Microscopy GmbH, Jena, Germany).

### Protein expression

GFP-Wash, Wash, GFP-Wash^ΔPav/Tum^, Wash^ΔPav/Tum^, Wash^ΔSHRC^, Wash^ΔArp2/3^ were amplified as 5’BamHI-3’NotI fragments by PCR and then cloned into a double-tag pGEX vector (GST and His; (Liu et al., 2009)). Primers used for cloning are described in Table S1. Protein expression assays were performed as previously described (Rosales-Nieves et al., 2006). Pav, PavDEAD, Tum, and CapuFH2 protein purification was performed as previously described (Nakamura et al., 2020; Rosales-Nieves et al., 2006). For Wash proteins, cells were lysed by sonication in T100G5 buffer (50 mM Tris, pH 7.6, 100mM NaCl, 5% Glycerol, and 1 mM DTT) with 1% Triton X-100, 50 mM imidazole, and complete protease inhibitor tablets (Roche).

Lysates were centrifuged at 10,000 g for 30 min, and the supernatants were coupled to Fastflow Nickel-Sepharose (GE) for 3 h at 4°C. The matrix was washed three times with lysis buffer with 50 mM imidazole and eluted by lysis buffer with 1 M imidazole. All His elutions, protein, were coupled to glutathione-sepharose 4B (GE) for 3 h at 4°C, washed with lysis buffer. PreScission protease (GE) was used to elute Wash proteins instead of reduced glutathione. All Wash proteins were dialyzed into storage buffer (50 mM Tris [pH 7.6], 50 mM NaCl, 1 mM DTT, 5% glycerol) and then flash frozen.

### F-actin/Microtubule (MT) bundling and cross-linking assays

Rabbit muscle actin (Cytoskeleton) was polymerized in polymerization buffer (10 mM Tris, pH 7.5, 50 mM KCl, 2 mM MgCl_2_, and 1 mM ATP) and then stabilized with Alexa Fluor 488 (or 633 for GFP-Wash and GFP-Wash^ΔPav/Tum^) conjugated Phalloidin. MT was polymerized by mixing unlabeled bovine brain tubulin (Cytoskeleton) and rhodamine-tubulin (Cytoskeleton) in a ratio of 1:5 and then stabilizing with paclitaxel (Cytoskeleton). MTs and test proteins were incubated in binding buffer A (80 mM Pipes, pH 7.0, 1 mM MgCl_2_, 1 mM EGTA) with 2 μM paclitaxel and 4 U/100 μl Alexa Fluor 488 (or 633) conjugated Phalloidin for 15 min at room temperature. F-actin was then added and incubated for 10 min. The mixture of protein, actin, and MTs was pipetted onto slides and then visualized by confocal fluorescence microscopy. Final protein concentrations for bundling assays: Wash^WT^, 500 nM; Wash^ΔSHRC^, 500 nM; Wash^ΔArp2/3^, 500 nM; Wash^ΔPav/Tum^, 500 nM; CapuFH2, 500 nM; sfGFP-Wash, 50 nM; and sfGFP-Wash^ΔPav/Tum^, 50 nM.

### Statistical analysis

All statistical analyses were done using Prism 8.2.1 (GraphPad, San Diego, CA). All graphs were generated using R-3.6.1. Statistical significance was calculated using a Two-tailed student’s t-test or a Kruskal–Wallis test for independence.

## Acknowledgements

We thank David Glover (Caltech), Li Tao (Univ. Hawaii), Paul Fisher (Stony Brook), Justin Hui and Toshi Tsukiyama (Fred Hutch), the Fred Hutch Proteomics, Antibody Development, and Cellular Imaging Shared Resources, Fred Hutch/Leica Center of Excellence, the Bloomington Stock Center, the Kyoto Drosophila Stock Center, FlyBase, and the Developmental Studies Hybridoma Bank, for advice, microscopes, antibodies, DNAs, flies, and other reagents used in this study.

## Competing Interests

The authors declare no competing or financial interests.

## Author Contributions

All authors performed experiments, contributed to the design and interpretation of the experiments, and to the writing of the manuscript.

## Funding

This research was supported by NIH GM143186 and the Mark Groudine Chair for Outstanding Achievements in Science and Service (to SMP), a Koss family donation to the Fred Hutch Basic Sciences Division, and NCI Cancer Center Support Grant P30 CA015704 (pilot project; Cellular Imaging, Antibody Development, and Proteomics Shared Resources).

## SUPPLEMENTARY FIGURE LEGENDS

**Fig. S1.**
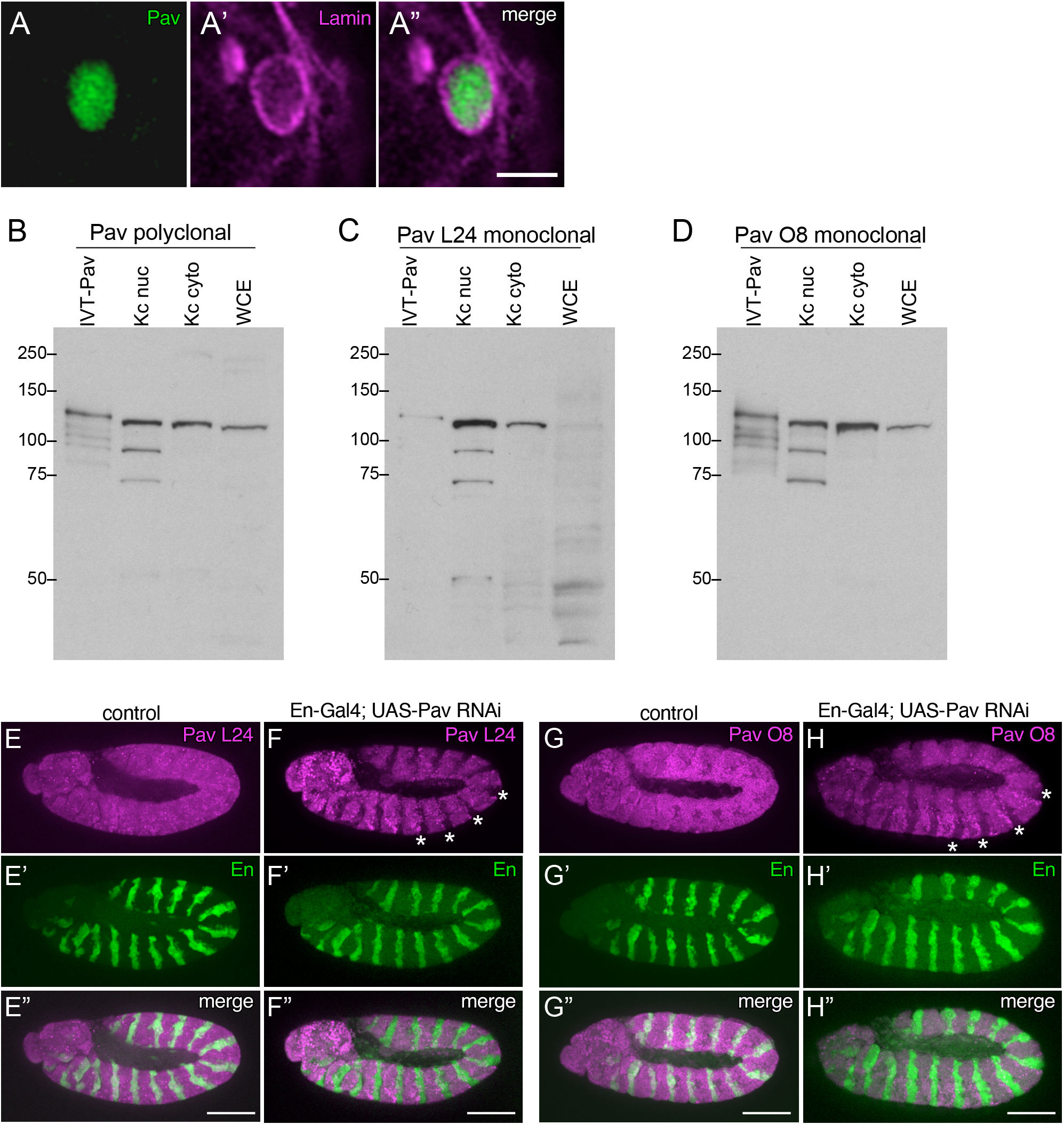
Pav antibody specificity. (A-A”) Super-resolution (Airyscan) micrograph projection of wildtype larval salivary gland nucleus stained with antibodies to Lamin and Pav. This rabbit polyclonal Pav antibody is from the Glover lab and described in Adams *et al*. 1998. Scale bar: 0.5μm. (B-D) Specificity of Pav mouse polyclonal and monoclonal lines L24 and O8. (B) Western blot of *in vitro* translated full-length Pav protein, Kc cell nuclear extract, Kc cell cytoplasmic extract, and whole cell 0-2hr Drosophila embryo extract probed with Pav polyclonal (B) or Pav monoclonal lines L24 (C) or O8 (D). (E-H”) Staining of control (E-E”, G-G”) or En-Gal4; UAS-Pav RNAi (F-F”, H-H”) embryos with antibodies to the Pav L24 (E-F”) or Pav O8 (G-H”) monoclonal lines to show antibody specificity. Note that these monoclonal antibodies do not stain the En regions where Pav is knocked down (asterisks). Scale bars: 100μm.

**Fig. S2.**
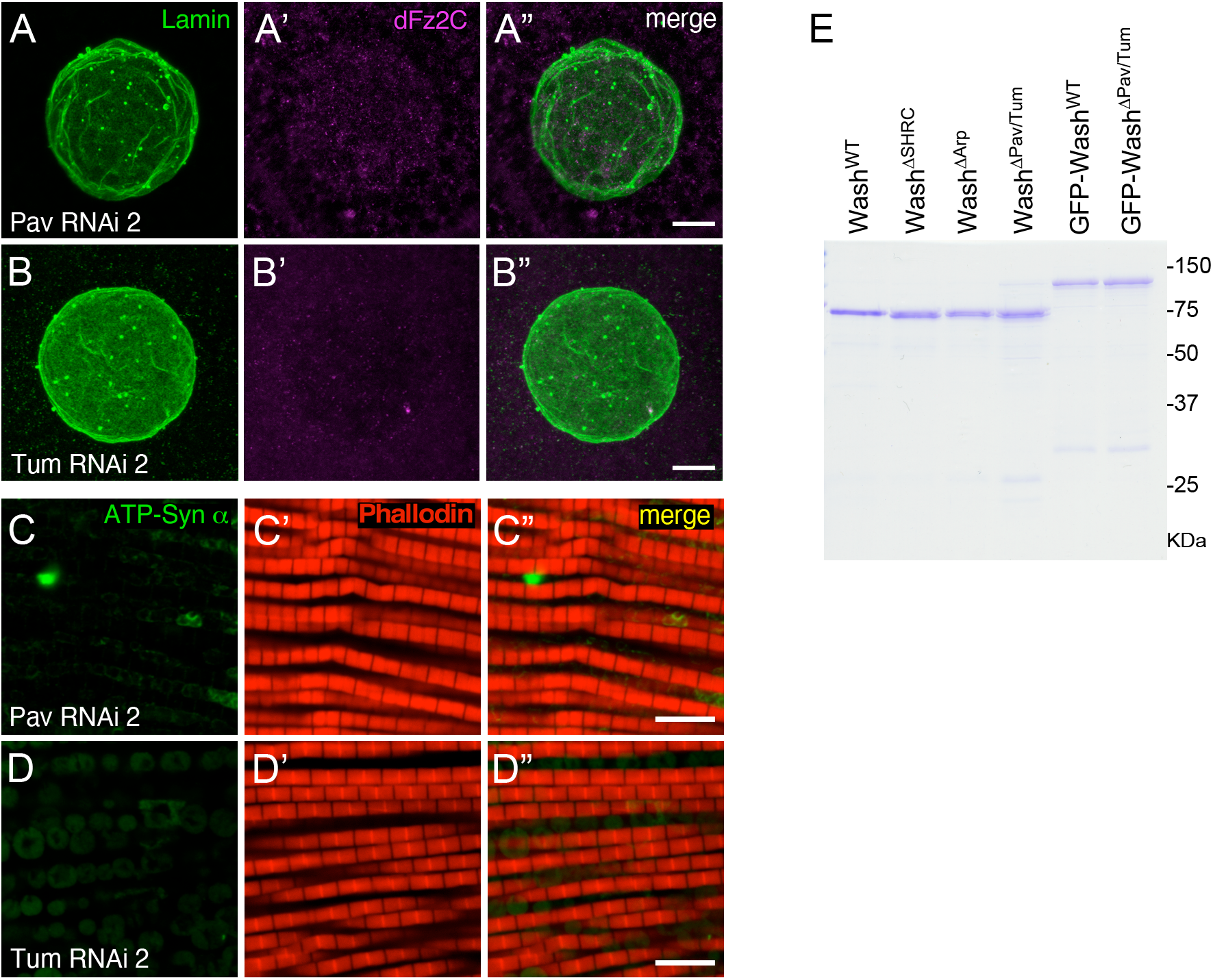
Pav/Tum mutant characterization. (A-B”) Confocal micrograph projections of Pav RNAi 2 (A-A”) and Tum RNAi 2 (B-B”) larval salivary gland nuclei stained with Lamin B and dFz2C. (C-D”) Confocal micrograph projections of adult IFM from Pav RNAi 2 (C-C”) and Tum RNAi 2 (D-D”) flies aged 21 days then stained for the activity dependent mitochondrial marker ATP-Syn α and Phalloidin. (E) Coomassie stained gel of bacterially purified proteins used for the F-actin bundling assays. Scale bars: 5μm.

**Table S1.**
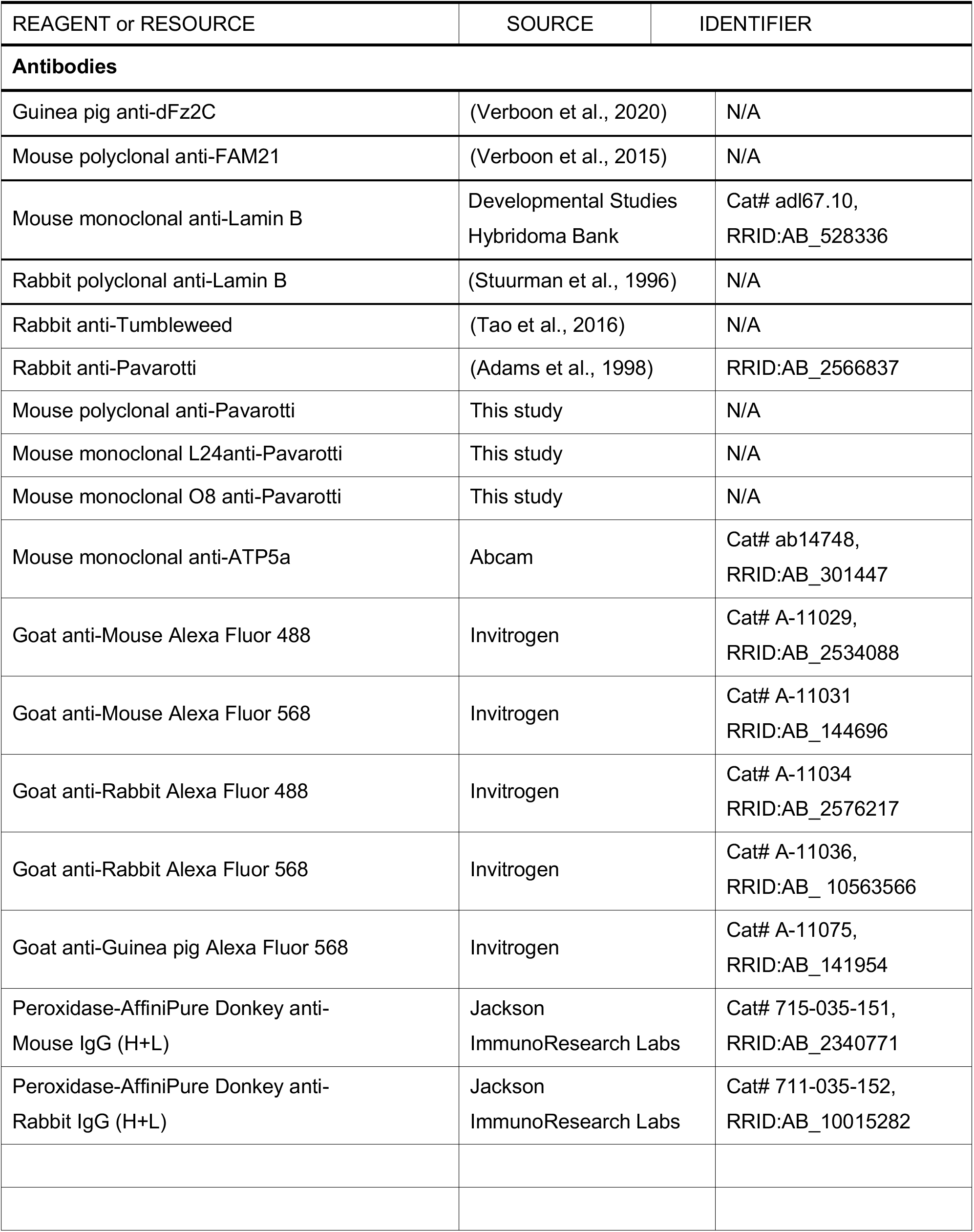
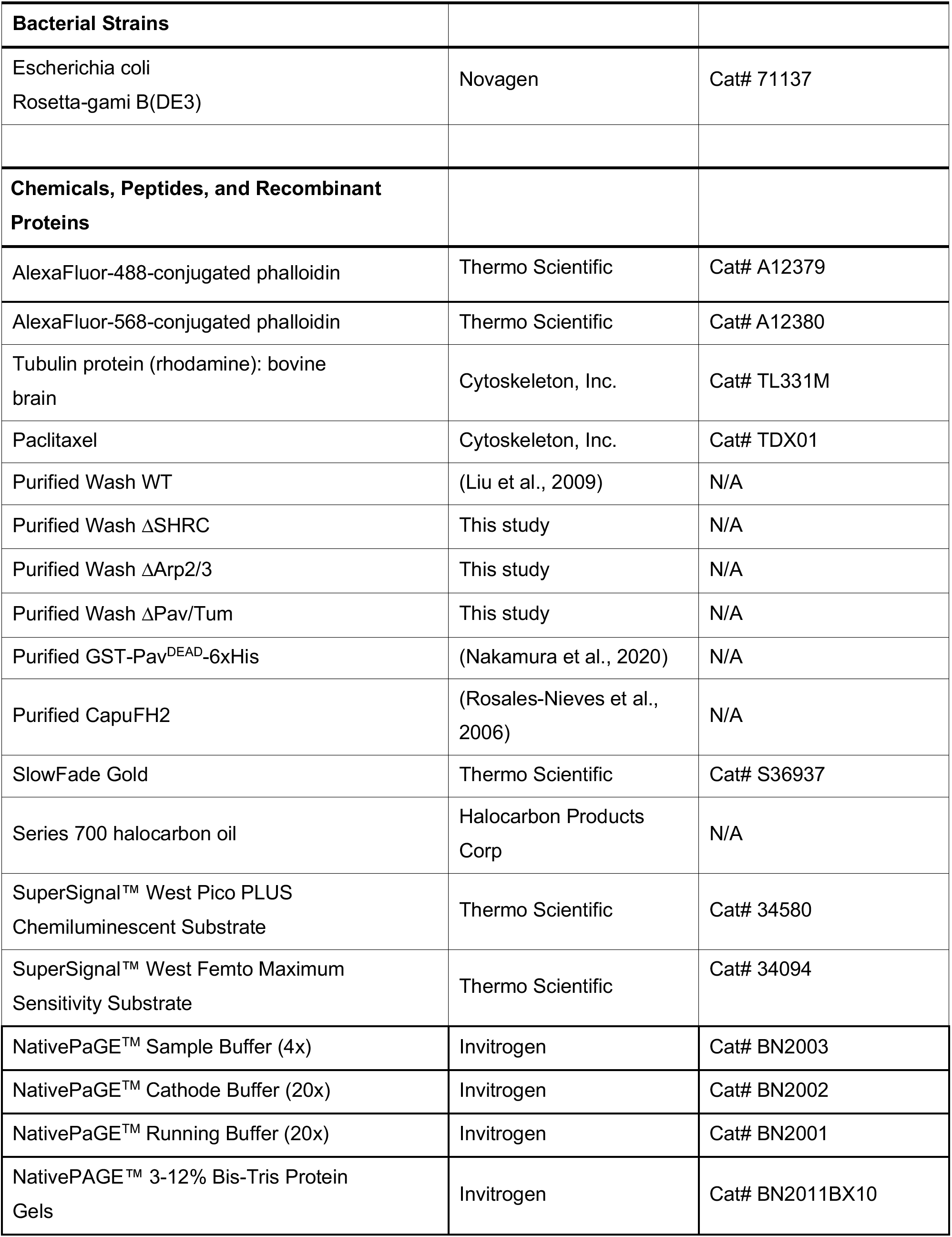
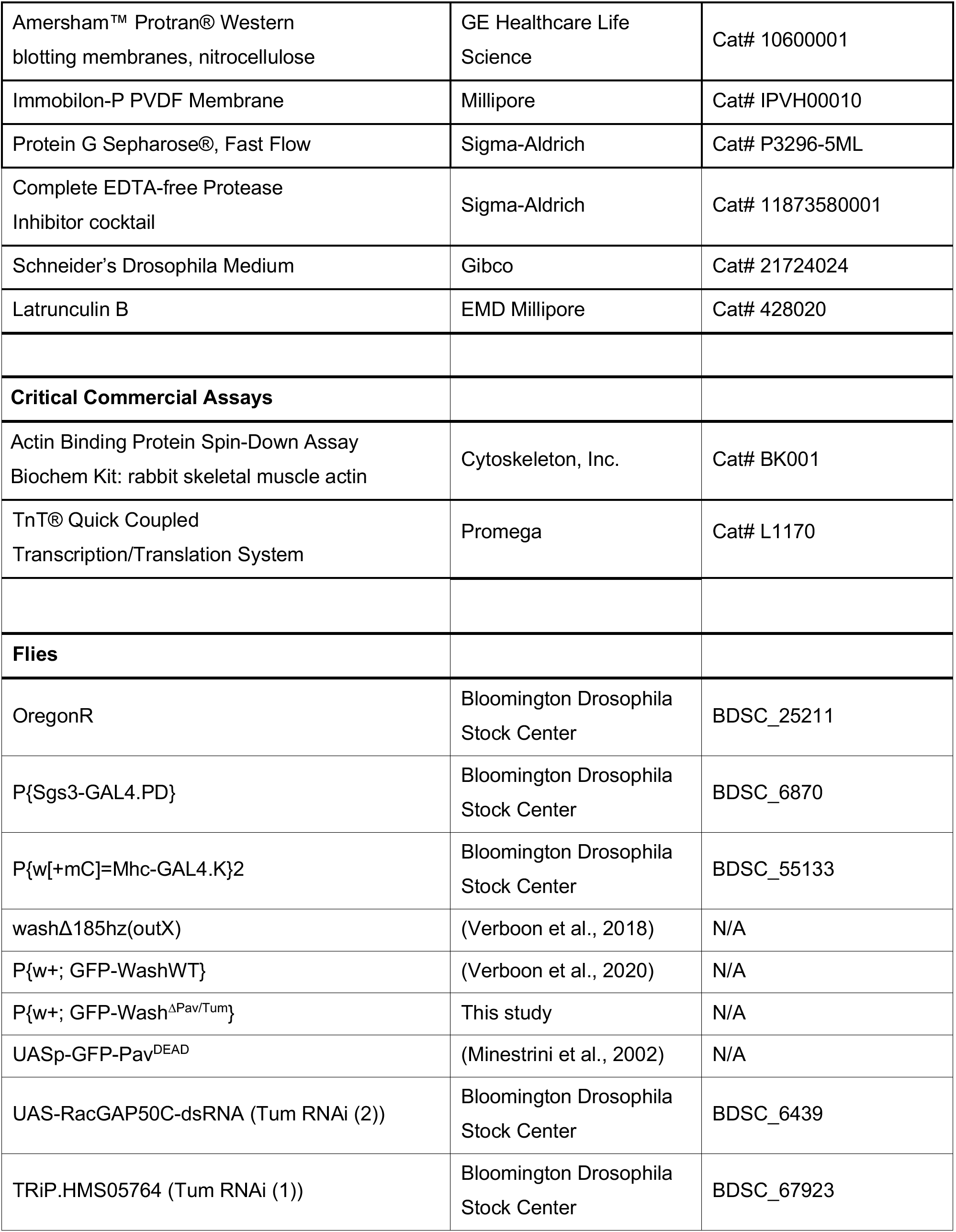
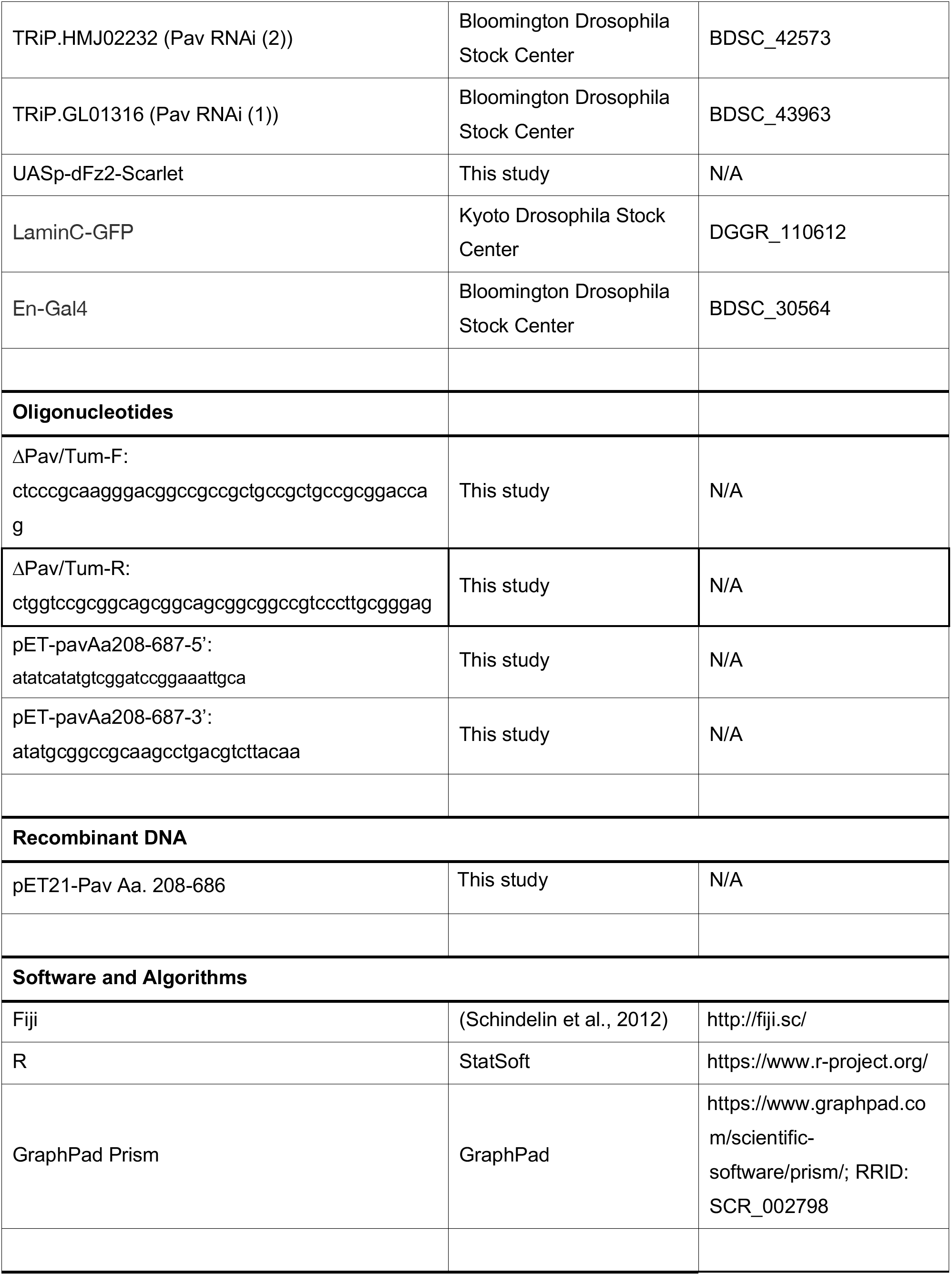
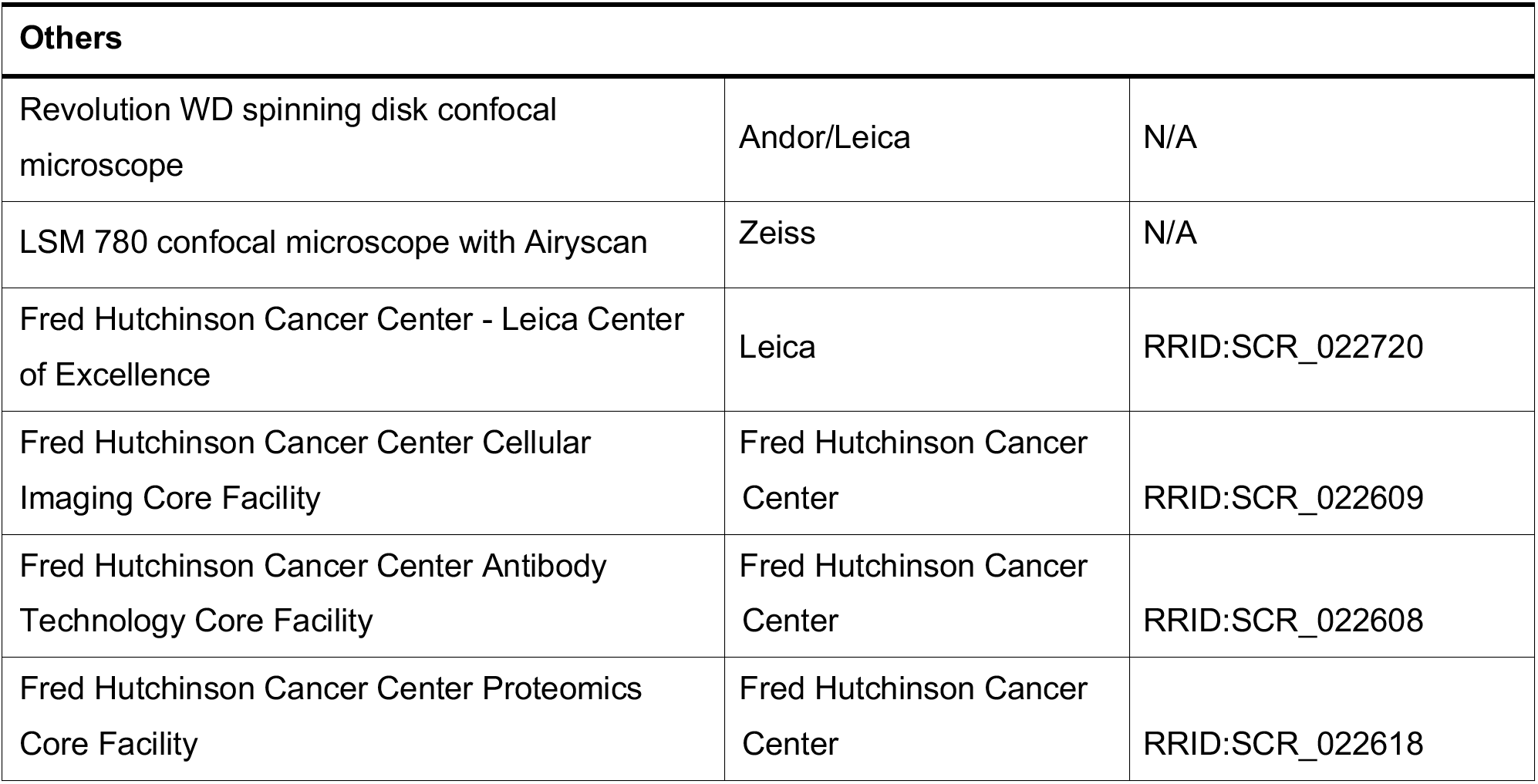
Reagents/Resources used in this study.

## Notes

### Competing Interest Statement

The authors have declared no competing interest.

